# Anti-CRISPR-mediated continuous directed evolution of CRISPR-Cas9 in human cells

**DOI:** 10.64898/2025.12.11.693673

**Authors:** Andrew L. Sabol, Amanuella A. Mengiste, Prashant Singh, Vedagopuram Sreekanth, Samuel J. Hendel, Minh Thuan Nguyen Tran, Anton M. Barybin, Santosh Chaudhary, Ra’Mal M. Harris, Kristi Liivak, Zachary C. Severance, Cale M. Locicero, Karishma Kailass, Chaiheon Lee, Lucy Qinghua Xu, Vincent L. Butty, Amit Choudhary, Matthew D. Shoulders

## Abstract

Engineering CRISPR-Cas systems for improved or altered function is central to both research and therapeutic applications. Unfortunately most optimization, especially directed evolution in bacterial hosts, fails to capture the functional requirements of the complex mammalian cellular milieu, where activity is usually required. Robust strategies to enable continuous directed evolution of genome-targeting agents directly in human cells remain lacking. Here, we introduce CRISPR-MACE (Mammalian cell-enabled Adenovirus-assisted Continuous Evolution) as a foundational technology to address this need. CRISPR-MACE integrates virus-based continuous evolution with anti-CRISPR-based tunable selection to generate novel *Streptococcus pyogenes* Cas9 variants with both increased and decreased DNA binding capacity and nearly 1000-fold–enhanced resistance to AcrIIA4, the strongest known inhibitor of SpCas9. Notably, across independent evolution campaigns the same Cas9 gatekeeper mutation reproducibly emerged first, enabling subsequent adaptive steps along two interdependent axes of Cas9 function. In addition to advancing CRISPR technologies, this work establishes key principles and synthetic circuits for continuously evolving CRISPR-Cas systems directly in human cells.

**SIGNIFICANCE STATEMENT:** CRISPR technologies are typically engineered in bacteria, even though they must function in the far more complex environment of human cells. This gap has limited the discovery of variants with improved DNA recognition or with resistance to inhibitors that operate differently in mammalian systems. Here we establish CRISPR-MACE, a continuous evolution platform that leverages pressure from anti-CRISPR proteins to select Cas9 variants directly in human cells that have novel functions. Evolved variants show improvements in DNA binding strength and residence time, as well as striking escape from the potent Cas9 inhibitor AcrIIA4. Many anti-CRISPR proteins use distinct mechanisms, so our strategy can drive future continuous evolution campaigns in mammalian cells that expand the functional properties of genome-targeting agents.

## INTRODUCTION

CRISPR-associated nucleases (e.g., Cas9) are RNA-guided endonucleases that recognize and neutralize bacteriophage DNA. The ease of targeting these nucleases to virtually any genomic locus is propelling the development of transformative CRISPR-based research tools and therapeutic agents.^1–3^ Over the past decade, numerous engineering approaches, especially directed evolution and computation-assisted rational design,^4–6^ have been employed to develop and optimize functionally diverse CRISPR-based genome targeting agents.^7–9^ Since CRISPR systems evolved as a bacterial defense mechanism against bacteriophages, the resulting reagents face the challenge that their activity in human cells is often sub-optimal. Indeed, several CRISPR-associated nucleases, while exhibiting potent activity in bacterial cells, are essentially inactive in human cells.^10, 11^ Such activity differences are understandable given the large differences in the size and structure of bacterial and human genomes, and the complex molecular recognition events (between DNA, RNA, and protein) that underlie the activity of these nucleases.

There is a growing consensus that directed evolution-based biomolecule engineering efforts should ideally be conducted in the same environments where biomolecular activity is desired.^12–32^ Unfortunately, efforts to evolve CRISPR-based functions directly in mammalian cells remain quite limited. Traditional step-wise discontinuous evolution approaches are difficult to implement at scale in human cells.^30, 31^ Although some mammalian cell-enabled targeted mutagenesis strategies have emerged,^20–29^ they are typically limited in mutational spectra. Moreover, their reliance on the cell itself as the unit of selection raises issues with selection escape and cheating. A fully integrated continuous evolution platform that, in analogy to the powerful phage-assisted continuous evolution (PACE) method that has been extensively applied to CRISPR system engineering in bacteria,^33^ enables simultaneous mutagenesis, selection, and amplification of CRISPR-based tools in human cells has yet to be achieved.

Here, we successfully develop such a platform and apply it to optimize the inhibitor-resistance and target DNA binding activities of Cas9, the most commonly used CRISPR-associated nuclease from *Streptococcus pyogenes*. The CRISPR-MACE platform reported here is a CRISPR-Cas tailored version of Mammalian cell-enabled Adenovirus-assisted Continuous Evolution.^13^ CRISPR-MACE co-opts adenovirus replication to enable continuous mutagenesis and selection of CRISPR systems. In CRISPR-MACE, a highly error-prone adenoviral DNA polymerase^34^ continuously generates Cas9 variants during adenoviral genome replication. Evolved Cas9 variants are then selected based on their ability to induce the expression of an essential, conditionally transcomplemented viral protease gene. We focused our selections on the DNA binding ability of Cas9. To help drive these selections, we relied on the anti-CRISPR proteins that bacteriophages themselves use to counter the bacteria’s CRISPR-based defense. The development and application of CRISPR-MACE reported herein paves the way for the use of continuous directed evolution in human cells to accelerate progress in the genome targeting field.

## RESULTS

### A tunable dCas9 selection circuit employing anti-CRISPR

CRISPR-MACE co-opts adenoviral replication to continuously perform all elements (mutagenesis, selection, and amplification) of a Cas9-focused directed evolution campaign directly in human cells. The platform employs an adenovirus that encodes a fusion of catalytically dead Cas9 with a VPR transcription activation domain (dCas9–TAD) but lacks two essential genes—adenoviral protease (AdProt) and adenoviral DNA polymerase (AdPol; **Figure 1a**). These genes are trans-complemented in engineered human cell lines able to inducibly express AdProt while constitutively expressing a highly error-prone variant of AdPol (EpPol)^34^ that introduces diverse mutations at rates similar to that of highly mutagenic RNA viruses throughout the viral genome (mutation rate of 3.7 × 10^-5^ for EpPol compared to ∼1.3 × 10^-7^ for AdPol),^13^ including in the dCas9–TAD gene. Notably, in CRISPR-MACE the human cell hosts merely provide the environment where virus-encoded dCas9–TAD variants are tested for function. The selector cells themselves are otherwise regularly discarded, promoting robust purifying selection and avoiding common cheating mechanisms that can plague continuous evolution campaigns in mutagenesis-only systems that rely on the cell as the unit of selection.

**Figure 1.**
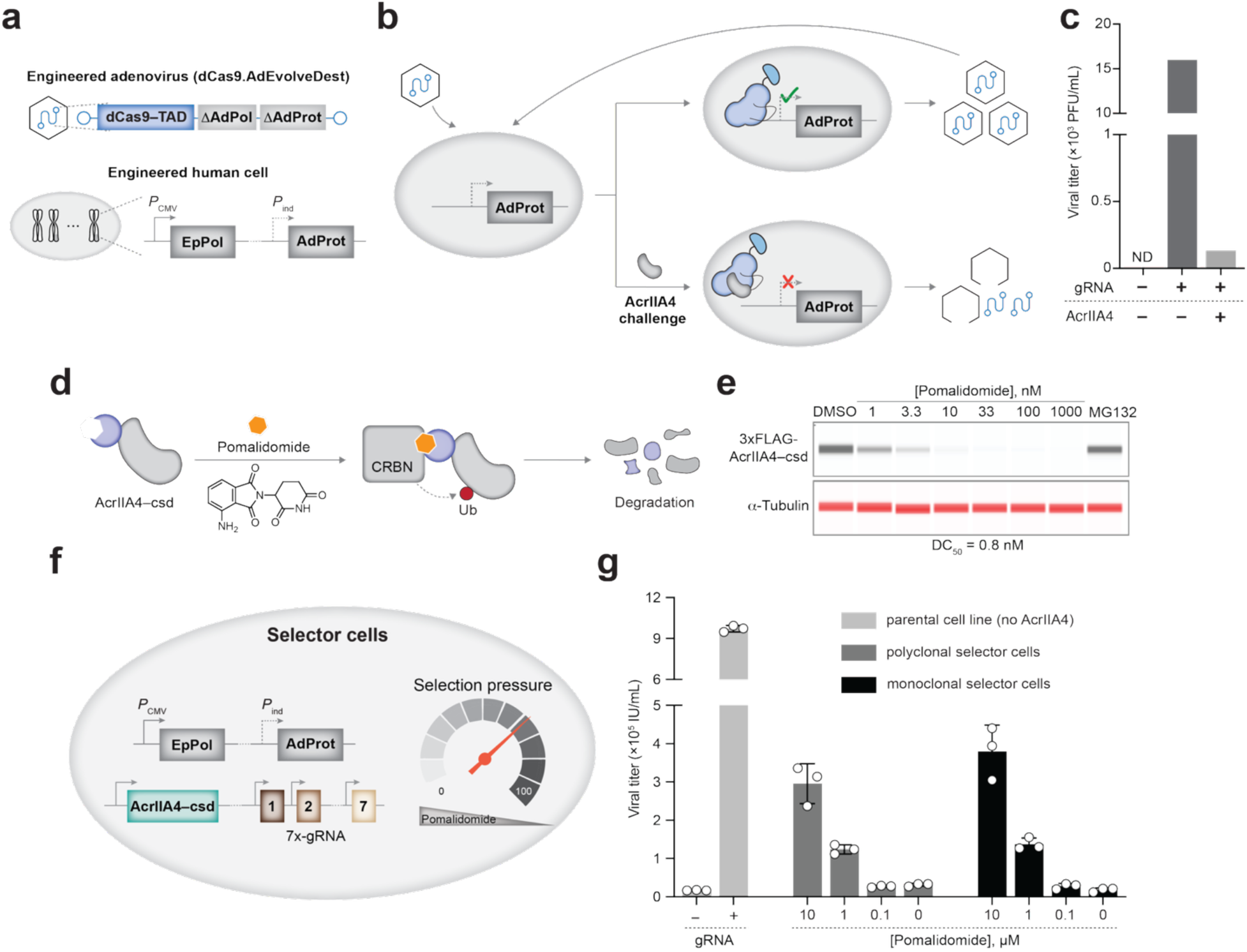
Establishment of CRISPR-MACE (<u>M</u>ammalian cell-enabled <u>Adenovirus</u>-assisted <u>C</u>ontinuous <u>E</u>volution). **(a)** CRISPR-MACE relies on a replication-deficient adenoviral genome (dCas9.AdEvolveDest) encoding dCas9–TAD and lacking both the adenoviral protease (AdProt) and DNA polymerase (AdPol). Adenoviral replication can occur only in complementing cells expressing error-prone AdPol (EpPol) and AdProt (note that the adenoviral genome is also lacking the E1 region, which is trans-complemented, and the E3 region that is not trans-complemented at all, thus ensuring that infection-competent adenoviruses cannot be generated in CRISPR-MACE). **(b)** EpPol is constitutively expressed from the host cells, while AdProt expression is activated by tiling gRNAs that target dCas9–TAD to the DNA upstream of the AdProt gene. AdProt production, and thus adenoviral replication, is suppressed by AcrIIA4. **(c)** Viral replication assay showing AdProt activation only in the presence of gRNAs and absence of AcrIIA4. Viral titers as plaque-forming units per mL (PFU/mL) were quantified. **(d)** Pomalidomide induces ubiquitination and proteasomal degradation of AcrIIA4 fused to a C-terminal superdegron (AcrIIA4–csd). **(e)** Capillary immunoblot showing dose-dependent degradation of AcrIIA4–csd, with MG132 stabilizing the protein as a positive control. **(f)** Schematic of selector cells stably encoding AcrIIA4–csd, a 7×gRNA cassette, EpPol, and AdProt. Pomalidomide levels fine-tune selection pressure in these cells by modulating AcrIIA4–csd abundance. **(g)** Viral replication assay demonstrating pomalidomide-dependent relief of selection pressure in selector cells. Viral titers as infectious units per mL (IU/mL) were measured by flow cytometry.

We configured CRISPR-MACE for Cas9 evolution by incorporating a promoter-less AdProt cassette into human cells such that, to produce the next generation of viable virions, viral genomes must encode and express variants of dCas9–TAD that can bind guide RNAs (gRNAs) targeting the region immediately 5′ of AdProt and successfully switch-on AdProt transcription (**Figure 1b**). In this manner, induction of AdProt transcription by dCas9–TAD results in the preferential selection and amplification of viral genomes carrying more active dCas9–TAD variants, which in turn infect new host cells and continuously undergo further rounds of evolution.

To apply selection pressure on DNA binding, we added the DNA-mimetic anti-CRISPR AcrIIA4 to the circuit (**Figure 1b**). Crucially, Cas9 recognizes target DNA by interacting with an upstream protospacer-adjacent motif (PAM) sequence, and AcrIIA4 is a competitive inhibitor of this Cas9-PAM binding interaction.^35–37^ Furthermore, AcrIIA4 also binds to the DNA unwinding and nuclease regions of Cas9.^35–37^ Thus, any dCas9–TAD variant that functions in the presence of AcrIIA4 must escape the DNA mimicry of AcrIIA4 while retaining (or even improving) the ability to bind human genomic DNA. We anticipated that this selection would present a unique evolutionary challenge, with unpredictable impacts on dCas9’s DNA-binding properties.

To assess the complexity of evolving functional dCas9 variants that are resistant to AcrIIA4, we evaluated the transcriptional activity of rationally designed Cas9 variants described in a past effort to elucidate the mechanism of AcrIIA4-mediated Cas9 inhibition.^35^ While the identified Cas9 variants were reported to suppress association with AcrIIA4, they additionally reduce or abolish transcriptional activity when evaluated as dCas9–TAD fusions (**Figures S1a, b**).

We experimentally validated our circuit by measuring the degree of viral replication under various conditions, including in the presence of AcrIIA4. We were able to robustly link adenoviral replication to dCas9–TAD function, as designed (**Figure 1c**). Moreover, co-expression of AcrIIA4 greatly reduced viral replication, indicating that there was a strong selective pressure for anti-CRISPR escape and altered DNA-binding (**Figure 1c**).

An initial, unsuccessful evolution campaign employing this circuit suggested to us that the selection pressure exerted by wild-type AcrIIA4 was insurmountably high during early viral passages. Indeed, as in **Figure 1c**, constitutive expression of wild-type AcrIIA4 effectively aborted viral propagation, preventing the emergence of any potential AcrIIA4 resistance-conferring substitutions in dCas9–TAD. While these observations reflect one valuable feature of our circuit—that it was not leaky—they also emphasized the need for a mechanism to fine-tune AcrIIA4 selection pressure.^38^ To address this need, we adopted a molecular glue-based degradation approach to titrate the levels of AcrIIA4.^39^ Briefly, we fused a 60-amino acid ‘superdegron’ element to the C-terminus of AcrIIA4 to create a fusion protein, AcrIIA4–csd, whose degradation could be driven by treatment with the small-molecule pomalidomide. The addition of pomalidomide induces proximity between AcrIIA4–csd and the cereblon E3 ligase, triggering AcrIIA4–csd ubiquitination and proteasomal degradation (**Figure 1d**).^39^ While AcrIIA4–csd alone strongly suppressed dCas9–TAD-driven AdProt transcription (a proxy for viral replication), this inhibition was substantially eliminated in AcrIIA4–csd-expressing cells upon pomalidomide treatment (**Figures S1c, d**). Crucially, the extent of AcrIIA4–csd degradation was proportional to the concentration of pomalidomide (**Figure 1e**), providing dose-dependent control over dCas9–TAD-mediated transcriptional activation (**Figures S1e, f**).

Given these promising results, we anticipated that pomalidomide dosing could, by tuning the amount of AcrIIA4–csd present in the cell at any given time, be used to modulate selection pressure on a dCas9–TAD-expressing adenovirus over the course of a directed evolution campaign. We used lentiviral transduction to stably integrate AcrIIA4–csd into a parental cell line that already genomically encoded EpPol (expressed from a constitutive promoter) and promoter-less AdProt. We were then able to derive a single monoclonal cell line that expressed AcrIIA4–csd as highly as during transient transfection (**Figure S2a**). As expected, dCas9–TAD-dependent viral replication was strongly inhibited in this monoclonal cell line in the absence of pomalidomide (**Figures S2b, c**). That inhibition was largely relieved by the addition of pomalidomide.

Next, we stably integrated a multi-gRNA cassette^40^ into this AcrIIA4–csd encoding monoclonal cell line to ensure that all elements necessary for our transcriptional activation selection circuit would be reliably maintained (**Figure 1f**). As shown in **Figure S2d**, gRNA-encoding monoclonal cell line #4—henceforth referred to as ‘monoclonal selector cells’—expressed high levels of several gRNAs, as did the gRNA-encoding polyclonal cell line (henceforth, ‘polyclonal selector cells’). Promisingly, genomic expression of gRNAs in both selection cell lines resulted in improved dCas9–TAD-dependent viral replication in the presence of pomalidomide compared to transient gRNA transfection (**Figure S2e**). Most importantly, the relief of selection pressure (enabled by pomalidomide-dependent AcrIIA4–csd degradation and measured as dCas9–TAD-dependent viral replication) was dose-responsive in both cell lines (**Figure 1g**).

### Robust enrichment of novel Cas9 variants in two independent evolution campaigns

With optimized se-lector cell lines in hand, we initiated two evolution campaigns by transducing either polyclonal (Campaign #1) or monoclonal (Campaign #2) selector cells with wild-type dCas9–TAD-encoding virus (**Figure 2a**). These campaigns enacted a pomalidomide-controlled selection pressure that filtered for viruses encoding variants of dCas9–TAD that escape AcrIIA4 inhibition while retaining their DNA binding capacity (**Figure 2b**). Proceeding with the campaigns simply involved occasionally passaging virus onto fresh selector cells while gradually decreasing the concentration of pomalidomide by dilution and thereby gradually increasing selection pressure (**Figure 2c**). The evolution campaigns were concluded when dCas9–TAD-encoding virus replicated robustly on selector cells at the highest applicable selection pressure—that is, in the absence of pomalidomide.

**Figure 2.**
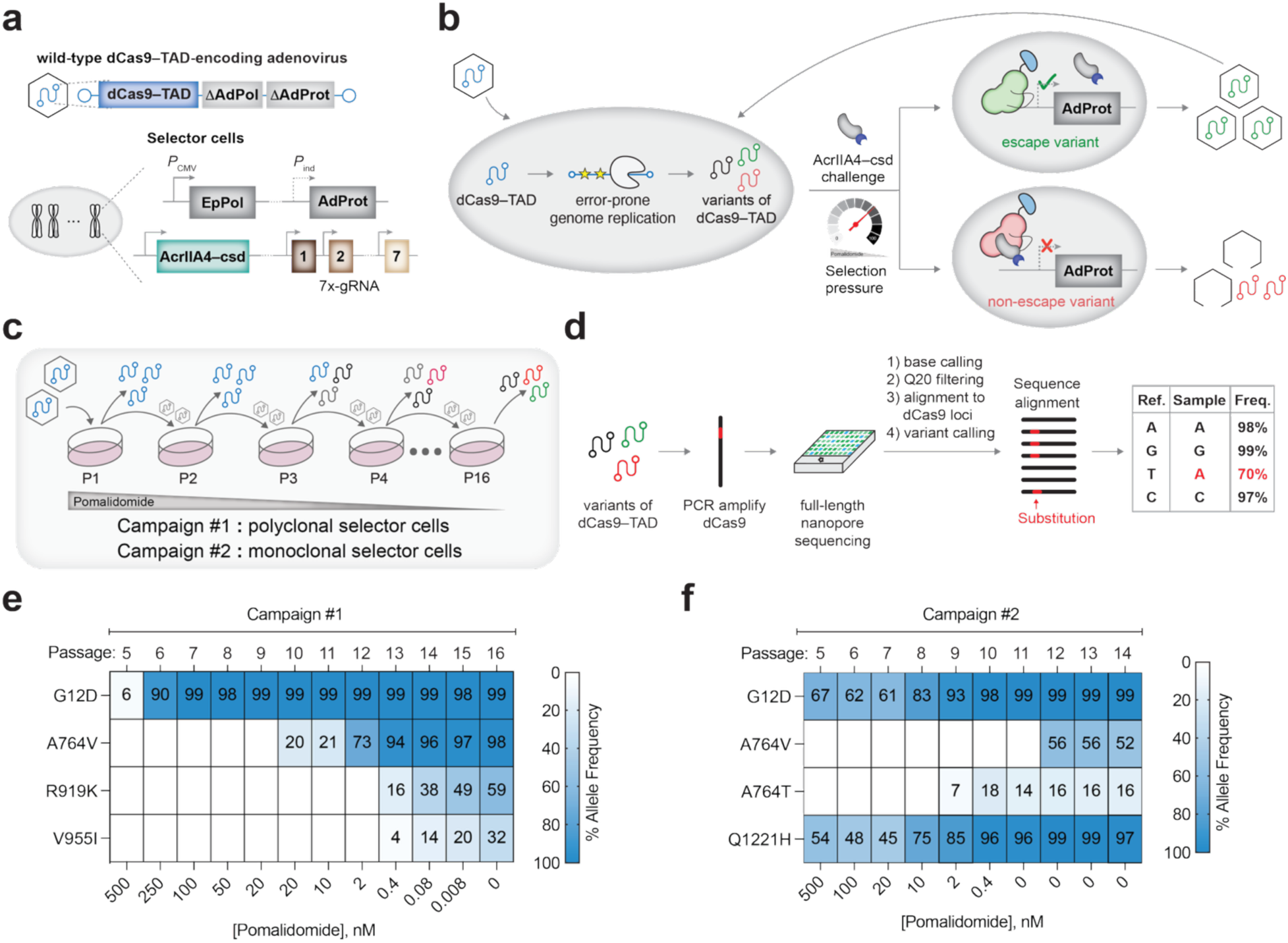
Directed evolution of dCas9–TAD using CRISPR-MACE. **(a)** Overview of components for the directed evolution of dCas9–TAD. Selector cells were infected with adenovirus encoding wild-type dCas9–TAD and lacking AdPol and AdProt. **(b)** In selector cells, constitutively expressed error-prone adenoviral polymerase (EpPol) introduces frequent mutations in dCas9–TAD. Selection pressure is tuned by pomalidomide-controlled degradation of AcrIIA4–csd. Variants that escape increasing amounts of AcrIIA4–csd inhibition and retain DNA binding activate AdProt expression, driving viral replication and enrichment. Non-escape or inactive variants result in reduced replication and elimination from the viral population. **(c)** Workflow for directed evolution in polyclonal (Campaign #1) and monoclonal (Campaign #2) selector lines. Virus from each passage was used to infect fresh selector cells, and viral DNA was sequenced to track evolutionary dynamics. **(d)** Nanopore sequencing pipeline for real-time monitoring of mutation emergence and allele frequency shifts. **(e, f)** Six dCas9 SNPs were strongly enriched in both campaigns. Notably, the G12D mutation independently emerged and became dominant early in both Campaigns #1 and #2.

We monitored the course of evolution campaigns by extracting viral DNA from culture medium following each passage, PCR-amplifying the dCas9 region, and then sequencing the resulting 4.1 kb DNA fragment using a MinION portable long-read sequencer from Oxford Nanopore Technologies (**Figures 2c, d**).^41^ Six different non-synonymous single nucleotide polymorphisms (SNPs) strongly enriched across both campaigns and were maintained through to the final viral passage, resulting in an equivalent number of unique substitutions at the amino acid level (**Figures 2e, f** and **Tables S1, S2**). Numerous other mutations also enriched to meaningful levels at intermediate passages (**Tables S1, S2**) but were ultimately not maintained in the population.

In both independent campaigns, the first mutation to enrich invoked a glycine-to-aspartic acid substitution at position 12 (G12D) of dCas9, which attained high allele frequency by Passage 6. Substitutions at position 764 (A764V and A764T) also independently occurred in Campaign #1 as well as Campaign #2, although in Campaign #2 they appeared in later passages (Passages 9–12). Substitutions R919K and V955I co-occurred during Passage 13 of Campaign #1 and steadily increased in frequency, while Q1221H co-occurred and enriched to fixation alongside G12D in Campaign #2. Of note, the robust enrichment of non-synonymous variants seen in our cam-paigns—indicative of strong purifying selection—is relatively rare among mammalian cell-based continuous evolution platforms reported to date.^13–16^

### Transcriptionally active dCas9 variants exhibit AcrIIA4 resistance in biochemical and cellular assays

We employed biolayer interferometry (BLI)^42, 43^ on recombinant proteins (**Figure S3**) to quantify the AcrIIA4 resistance of our evolved dCas9 variants. We used a competitive binding assay,^36, 44^ in which the association and dissociation kinetics of dCas9–gRNA ribonucleoprotein (RNP) complexes with target double-stranded DNA (dsDNA) were measured in the presence or absence of AcrIIA4 (**Figure 3a**). Briefly, dCas9–sgRNA complexes were pre-incubated with varying concentrations of AcrIIA4 and then allowed to bind to biotinylated dsDNA immobilized on streptavidin biosensors. In the absence of AcrIIA4, wild-type dCas9 displayed stable binding to the target DNA. Pre-incubation with increasing concentrations of AcrIIA4 caused a dose-dependent decrease in association rate, ultimately leading to complete inhibition of dsDNA binding (**Figure 3b**). The resulting binding curves were normalized to the “RNP without AcrIIA4” condition and plotted against AcrIIA4 concentration to calculate the half-maximal inhibitory concentration (IC_50_), which was determined to be 9.3 nM for wild-type dCas9 (**Figures 3c, d** and **Table S3**), consistent with a previous report.^36^

**Figure 3.**
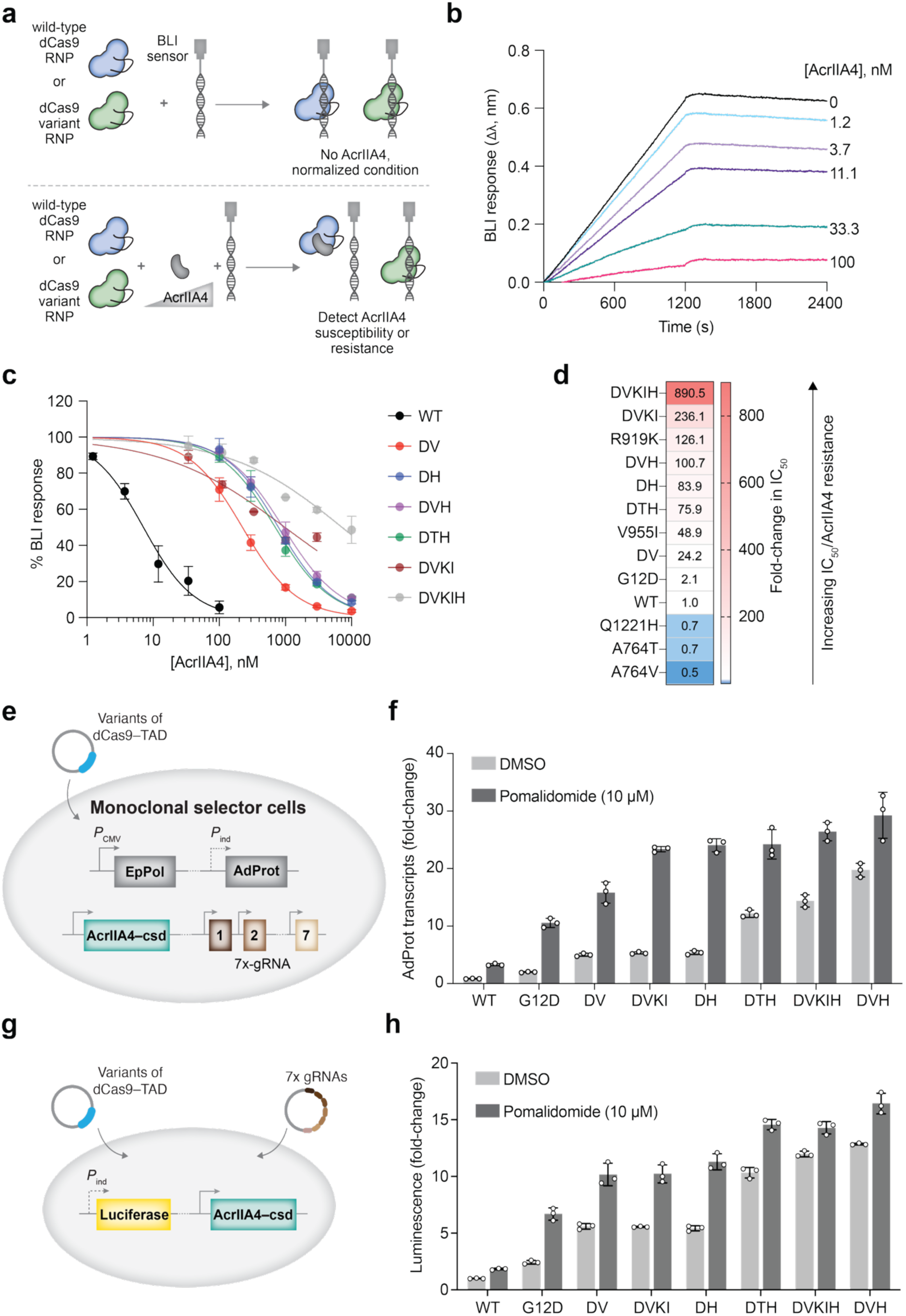
Biochemical and cellular characterization of AcrIIA4 resistance in evolved dCas9 variants. **(a)** Schematic of biolayer interferometry (BLI) assay measuring binding of wild-type or mutant dCas9 RNP complexes to immobilized dsDNA across AcrIIA4 concentrations. **(b)** Example of BLI response curves. Binding curves were normalized to the 0 nM AcrIIA4 condition and plotted against AcrIIA4 concentration to calculate half-maximal inhibitory concentration (IC_50_) **(c)** Binding (as % response) versus AcrIIA4 concentration plots for wild-type (WT) and multiply-substituted dCas9 variants. The inflection point of each curve equals the AcrIIA4 IC_50_ value corresponding to each dCas9 variant. **(d)** Heatmap of AcrIIA4 IC_50_ fold-changes relative to WT dCas9, indicating varying levels of AcrIIA4 resistance. **(e, f)** Monoclonal selector cells transfected with individual dCas9–TAD variants or a composite quintuple mutant (**e**) were treated with DMSO or pomalidomide, and AdProt transcript levels were quantified by qPCR (**f**). **(g, h)** HEK293T cells expressing nanoluciferase and AcrIIA4–csd were transfected with dCas9–TAD variants and luciferase-targeting gRNAs (**g**) to assess luciferase activation in the absence or presence of pomalidomide (**h**).

Using the aforementioned competition assay, we next recorded dose-response BLI curves and quantified the fold-change in AcrIIA4 IC_50_ for a series of dCas9 variants (**Figures 3c, d, S4, S5,** and **Table S3**). Among the single substitutions, G12D, which was the earliest variant to emerge in both campaigns, surprisingly conferred only a small increase in IC_50_, approximately 2-fold relative to wild-type dCas9. The Q1221H substitution, which appeared simultaneously with G12D in campaign #2, also did not alter IC_50_ significantly on its own. Neither did the A764V (or A764T) substitutions from both campaigns. Notably, the R919K (126-fold increase) and V955I (49-fold increase) substitutions, both of which only appeared later in the evolutionary trajectories, did provide substantial AcrIIA4 escape phenotypes as single substitutions.

We next examined the evolutionarily relevant combinations of these substitutions to determine how they cooperatively influence AcrIIA4 resistance. The double mutant G12D/A764V (DV) exhibited moderate resistance, increasing the IC_50_ by ∼25-fold, and the G12D/Q1221H (DH) combination yielded an ∼84-fold increase (**Figures 3c, d, S4, S5,** and **Table S3**). Resistance increased further in the G12D/A764T/Q1221H (DTH) and G12D/A764V/Q1221H (DVH) triple mutants, with IC_50_ values elevated by 75- to 100-fold, respectively. Strikingly, when the additional mutations R919K and V955I were introduced to the DV mutant to generate the quadruple mutant G12D/A764V/R919K/V955I (DVKI) observed in Campaign #1, AcrIIA4 resistance surged, reaching a 236-fold increase in IC_50_ relative to wild-type dCas9. The most dramatic effect was observed in the composite quintuple mutant G12D/A764V/R919K/V955I/Q1221H (DVKIH), which displayed nearly three orders of magnitude greater resistance to AcrIIA4 (with an 890-fold increase in IC_50_), effectively preventing inhibitor-mediated disruption of DNA binding even at high AcrIIA4 concentrations (**Figures 3c, d, S5f,** and **Table S3**). This progressive increase in IC_50_ values underscores the cooperative and synergistic effects of combining mutations and highlights how directed evolution can generate complex resistance profiles against potent natural inhibitors, such as AcrIIA4.

We next evaluated AcrIIA4 resistance and the retention of transcriptional activity for selected dCas9 variants in cells. Focusing on non-synonymous variants that were maintained at high levels even in the final passage, we cloned wild-type dCas9–TAD, along with the G12D, DH, DV, DTH, DVH, DVKI, and the composite DVKIH variants into a mammalian cell expression vector. This panel includes the single and multiple (1+) substitution variants that strongly enriched across each directed evolution campaign, as well as the DVKIH quintuple mutant that combines unique substitutions observed in separate campaigns. We transfected these constructs into monoclonal selector cells and used reverse transcription quantitative PCR (RT-qPCR) to assess whether the encoded dCas9–TAD variants had gained resistance to AcrIIA4–csd while maintaining transcriptional activity (**Figure 3e**).

We observed that all variants exhibited AcrIIA4–csd resistance in this cellular assay. The DVH triple mutant from Campaign #2 performed the best, while the DTH and composite DVKIH variants displayed similarly high resistance levels (**Figure 3f)**. Notably, the resistance improved as new (later passage) mutations were introduced. Moreover, resistance to AcrIIA4–csd was observed both in the presence and absence of pomalidomide. Reassuringly, we obtained similar results when a HEK293T cell line stably expressing a promoter-less luciferase alongside AcrIIA4–csd was transiently transfected with the same set of gRNAs now targeting the dCas9–TAD variants to the luciferase locus (**Figures 3g, h**). Thus, the improvement in resistance to AcrIIA4 observed was not specific to just one genomic locus.

We questioned whether the evolved dCas9–TAD variants had generally gained resistance to AcrIIA4, or just specifically gained resistance to the AcrIIA4–csd fusion protein. To address this question, we used cells stably expressing wild-type AcrIIA4 at levels similar to those attained in our AcrIIA4–csd-expressing selector cells in the absence of pomalidomide (i.e., in the absence of induced AcrIIA4–csd degradation; **Figure S6a**). Upon transient transfection of the full panel of dCas9–TAD variants into these wild-type AcrIIA4 expressing cells (**Figure S6b**), which also incorporated a promoter-less AdProt cassette and AdProt-targeting gRNAs, we observed AdProt induction trends (**Figure S6c**) recapitulating those observed in **Figures 3f** and **3h**. Overall, these biochemical and cellular assays exhaustively validate the capacity of CRISPR-MACE to yield transcriptionally active, AcrIIA4-resistant dCas9 variants. Notably, in contrast to past rational design efforts for AcrIIA4 resistance (**Figures S1a, b**), top-performing dCas9–TAD variants from our evolutionary campaigns, including the DVKIH, DVH, and DH variants, all retained wild-type or better levels of transcriptional activity in the absence of AcrIIA4 (**Figures S6d, e**).

### dCas9 variants have altered DNA-binding properties in biochemical and cellular assays

Next, we used a BLI assay to evaluate anticipated changes in the DNA binding affinity and kinetics of evolved dCas9 variants. Varying concentrations of dCas9–gRNA RNP complexes, including the benchmark wild-type Cas9, were incubated with biotinylated dsDNA immobilized on streptavidin biosensors (**Figure 4a**). BLI response curves were fitted using a 1:1 binding model, allowing us to compute the association rate constant (*k*_on_), dissociation rate constant (*k*_off_), residence time (τ), and binding constant (*K*_d_) for each variant (**Figures 4b–d**, **S7**, and **Table S4**). A dsDNA substrate lacking a PAM exhibited negligible binding (**Figures S8** and **S9**). For wild-type dCas9, the *k*_on_ and *k*_off_ values were 0.8 × 10⁴ M⁻¹s⁻¹ and 5.4 × 10⁻⁵ s⁻¹, respectively, yielding a *K*_d_ of 7.1 ± 0.7 nM (**Figures 4c, d**, **S7**, and **Table S4**), which is in agreement with a previous report.^42^ We then determined the *K*_d_ and τ of all evolutionarily relevant dCas9 variants, and identified some variants with 10-fold enhanced DNA binding (**Figures 4c, d, S7**, and **Table S4**). Conversely, some variants exhibited weaker DNA binding and decreased residence times.

**Figure 4.**
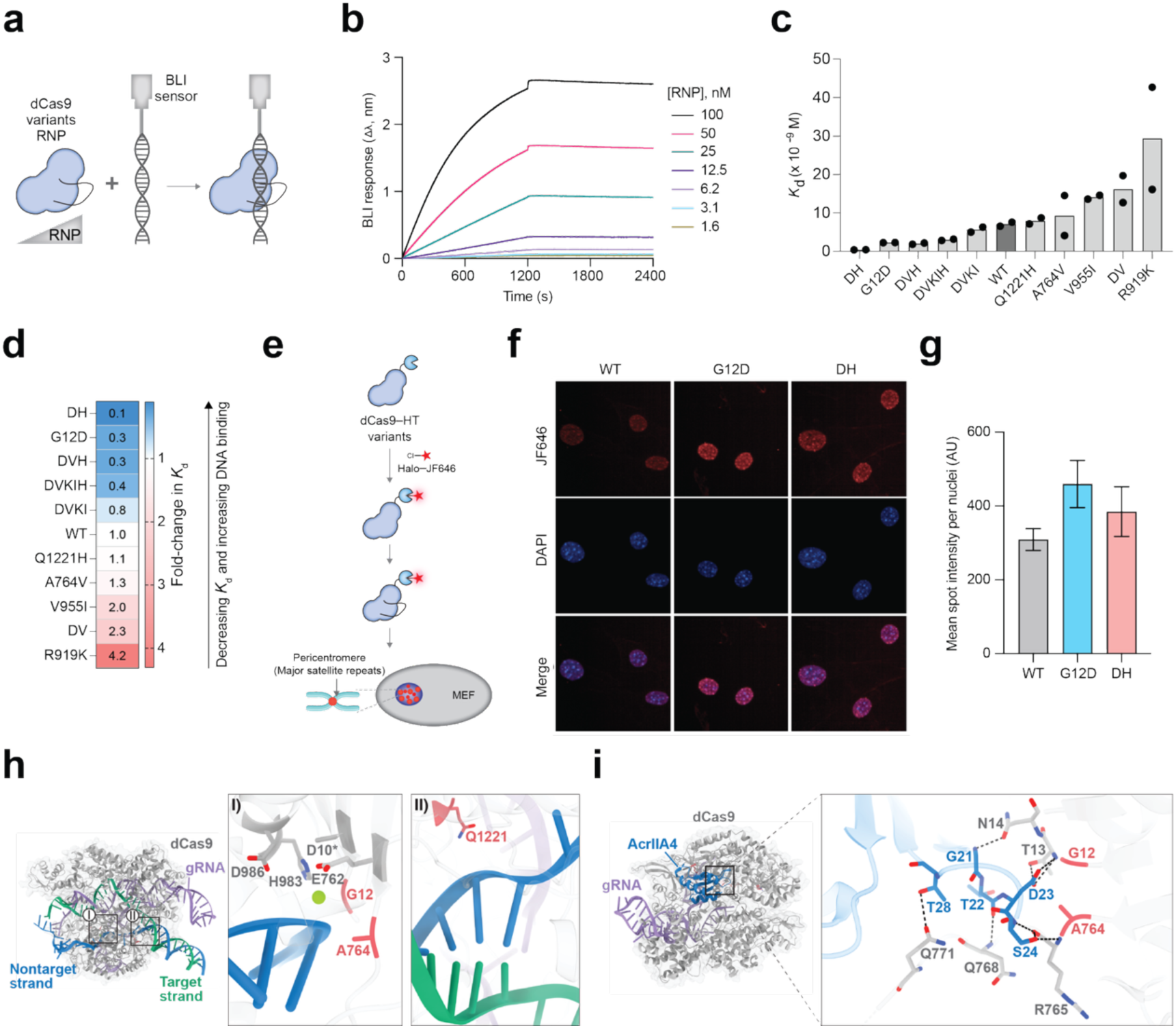
Biochemical and cellular characterization of DNA binding by evolved dCas9 variants. **(a)** Schematic of biolayer interferometry (BLI) assays measuring binding affinities of evolved dCas9 variants across RNP concentrations. **(b)** Representative binding curves for wild-type dCas9 (WT) with PAM-containing dsDNA; kinetic parameters (*k*_on_, *k*_off_, *K*_d_) were derived from dose–kinetic analysis. **(c, d)** Bar graphs of *K*_d_ values showing the effects of dCas9 substitutions on DNA-binding affinity (**c**) and heatmap representation of *K*_d_ fold-changes relative to WT dCas9 (**d**)**. (e)** Schematic of dCas9–HaloTag (HT) labeling and imaging experiment. dCas9–HT conjugated to JF646 was assembled into RNPs targeting pericentromeric repeats and imaged in fixed murine embryonic fibroblasts (MEFs) by confocal microscopy. **(f, g)** Confocal images of MEF nuclei transfected with WT, G12D, or G12D/Q122H (DH) dCas9–HT–JF646 RNPs (**f**), and quantification of satellite foci in >10,000 cells (**g**) showing variant-dependent DNA binding. **(h)** Locations of substituted residues in the Cas9–gRNA–dsDNA ternary complex (PDB 5Y36).^45^ Mutated residues (*red*) are shown relative to the RuvC domain and bound DNA. **(i)** Locations of substituted residues in the Cas9–gRNA–AcrIIA4 complex (PDB 5VW1),^37^ highlighting their positions near the AcrIIA4 interaction interface and associated hydrogen bonds (*black dashed lines*).

We analyzed the effects of individual substitutions beginning with G12D, the earliest mutation to emerge in both evolution campaigns. This substitution considerably enhanced DNA binding by lowering the *K*_d_ to 2.3 nM—a 3-fold improvement over wild-type dCas9 (**Figures 4c**, **d**, and **Table S4**). This variant also had a 1.8-fold improvement in residence time relative to wild-type dCas9 (**Figure S7c** and **Table S4**). The Q1221H substitution, which co-enriched with G12D in campaign #2 and was consistently present in the highest-performing cell-based variants, had little effect on its own (*K*_d_ = 7.9 ± 1.1 nM; **Figures 4c**, **d, S7** and **Table S4**). However, when combined with G12D to generate the DH double mutant, DNA binding was dramatically improved, displaying a ∼10-fold increase in DNA affinity relative to wild-type dCas9 (*K*_d_ = 0.4 nM; **Figures 4c**, **d, S7** and **Table S4**). This observation indicates a strong synergistic effect between these two substitutions in enhancing DNA target engagement.

Given their increased DNA binding affinities in biochemical settings, we hypothesized that the G12D and DH variants would be useful for *in cellula* applications where Cas9’s residence on DNA is crucial. To test this hypothesis, we employed these variants in a Cas9-mediated fluorescence *in situ* hybridization (CASFISH) assay, which uses an *in vitro* assembled fluorescently-labeled RNP complex to detect and image genomic loci.^46^ To this end, we assembled CASFISH probes by forming RNP complexes between purified Janelia fluorophore 646-labeled dCas9 variants (**Figure S10**) and gRNA. The resulting RNP complexes targeted the major satellite repeats of murine pericentromeres, a key chromosomal region for maintaining chromosomal integrity and physical spacing between centromeric (spindle attachment point in mitosis) and euchromatic chromosomal regions (**Figure 4e**).^47^ We observed nuclear-localized fluorescent punctate signals, congruous with pericentromeric regions, and detected higher punctate fluorescence intensity in the G12D and DH variants compared to wild-type dCas9 (**Figures 4f, g**), consistent with their greater DNA binding affinity.

In contrast to the G12D and DH variants, several other substitutions critical for AcrIIA4 resistance—including A764V, R919K, and V955I—strikingly weakened DNA binding when introduced individually. The A764V variant displayed a *K*_d_ of 9.3 ± 7.3 nM, while V955I and R919K were further impaired, exhibiting *K*_d_ values of 14.1 ± 0.7 nM and 29.4 ± 18.8 nM, respectively (**Figures 4c**, **d**, **S7**, and **Table S4**). These reductions suggest that, while these variants are quite beneficial for AcrIIA4 resistance (for example, the DV variant displays weaker DNA binding but higher transcriptional activity in selector cells with residual AcrIIA4 expression, as shown in **Figures 3f** and **3h**), they partly compromise intrinsic DNA binding capability. Importantly, when these substitutions were combined with the high DNA affinity G12D and DH backgrounds, the reductions in DNA binding capacity are partially offset, leading to intermediate *K*_d_ values. Thus, the fully evolved composite DVKIH variant retained robust DNA binding while achieving maximal AcrIIA4 resistance. Indeed, this variant showed slightly improved DNA affinity relative to wild-type dCas9, with a *K*_d_ of 3.0 ± 0.2 nM (**Figures 4c**, **d**, **S7**, and **Table S4**).

Based on these observations, G12D appears to be a key “gatekeeper” mutation, conferring improved DNA affinity that opens the door to access AcrIIA4-resistant alleles. G12D occurred first in both independent evolution campaigns, despite minimally improving AcrIIA4 resistance (**Figure 3d**). The Q1221H substitution that emerged alongside G12D in Campaign #2 similarly did not meaningfully confer AcrIIA4 resistance on its own, but improved DNA affinity when combined with G12D. The G12D and Q122H substitutions then served as a foundation upon which beneficial substitutions that more strongly conferred AcrIIA4 resistance could arise, despite their otherwise deleterious effects on DNA binding (**Figures 4c, d**). These results reveal a complex evolutionary landscape for dCas9–TAD evolution in CRISPR-MACE, with selection on two interdependent functional axes (DNA binding improvement versus AcrIIA4 binding reduction). Both historical contingency and epistatic effects appear to play key roles in the dCas9–TAD variants that ultimately emerged.

## DISCUSSION

Recognition of the importance of engineering biomolecular function directly in the environment where a given biomolecule is ultimately intended to be used has catalyzed the recent emergence of several new mammalian cell-based continuous directed evolution platforms that fully integrate all steps of a directed evolution campaign.^13–16^ Yet, none of these methods have been applied to one of the most impactful types of biomolecules—genome targeting agents like CRISPR-Cas proteins. Here, we developed and applied CRISPR-MACE to evolve the DNA binding properties of dCas9. We uncovered novel dCas9 sequences with both altered DNA affinity and remarkable resistance to the anti-CRISPR protein AcrIIA4. Moreover, multiple campaigns revealed a complex evolutionary landscape, consistent with a selection circuit mimicking the evolutionary molecular warfare between bacterial and bacteriophage proteins, but here played out in the human cell milieu.

Our evolution campaigns identified dCas9 variants bearing multiple mutations, which altered their DNA binding properties and substantially increased their resistance to AcrIIA4. We identified variants that exhibited a 10-fold increase in DNA binding affinity (and residence time) while others showed a 4-fold decrease. We then employed variants with enhanced DNA binding affinity to improve DNA imaging sensitivity at pericentromeres by 2-fold. We also identified variants of dCas9 that exhibited tens- to hundreds-fold resistance to AcrIIA4. Notably, many evolved variants are novel and mutations span multiple Cas9 domains.

The impact of some of the evolutionarily identified substitutions can be structurally rationalized. For instance, residues G12 and A764—located near the RuvC catalytic core of Cas9—engage in extensive interactions with both AcrIIA4 and the protospacer adjacent motif (PAM)-bearing DNA strand. The glycine-to-aspartic acid G12D gatekeeper substitution observed first in both directed evolution campaigns may enable Mg^2+^ ion coordination and thereby restore or partially recapitulate the hydrogen bonding network within the RuvC catalytic core that is abrogated by the nuclease-inactivating D10A substitution in dCas9 (**Figure 4h**), resulting in this key variant’s enhanced DNA affinity. On the other hand, the alanine-to-valine substitution at position 764 may weaken DNA affinity by introducing steric repulsion within the RuvC domain; a phenomenon that is likely exacerbated by the incorporation of a bulkier amino acid (G *versus* D) at position 12 (**Figure 4h**). Because A764 resides at the protein–protein interface formed by the association of AcrIIA4 and Cas9, the installation of larger amino acids at these loci likely sterically obstructs that association (**Figure 4i**) and is the basis of the AcrIIA4 resistance that substitutions at A764 confer. The impact of the Q1221H substitution can be similarly structurally rationalized given that residue Q1221—located in the PAM-interacting (PI) domain of Cas9—has a crucial interaction with the phosphate backbone of the non-target strand (**Figure 4h**). Indeed, the asparagine-to-histidine substitution observed at this residue was previously reported in a PAM-relaxed variant of Cas9, where it was proposed to improve binding of the non-target strand.^48^ Notably, we found that Q1221H can remarkably improve DNA affinity, but only if introduced alongside the G12D substitution. The origin of this epistatic effect is currently not clear. Curiously, although the R919K and V955I substitutions (both in the RuvC domain) substantially reduce AcrIIA4-Cas9 association, these residues are positioned more than 10 Å away from any Cas9:AcrIIA4 interface (**Figure S11a**), and residue 955 is distal to all Cas9:DNA binding interfaces (**Figure S11b**). The enrichment of these highly beneficial substitutions that appear to lack structural explicability, along with the requirement of the G12D gatekeeper substitution for their beneficial effects to emerge, typifies a long-established advantage of directed evolution—the identification of improved biomolecular variants and variant combinations that would otherwise elude rational analysis.^5, 49^

The successful outcomes of these directed evolution campaigns relied on several key features of CRISPR-MACE:

1. The high error rate of EpPol and continuous viral growth, which impart a large library size and diversity during *in cellula* mutagenesis, while avoiding the requirement for library generation via error-prone PCR. Notably, this mutagenesis underpins the robust generation and selection of multiply mutated variants.
2. The use of stably integrated synthetic circuits that avoid leakiness and provide consistent results across directed evolution campaigns.
3. A robust selection. Because the human cells merely serve as constantly discarded temporary hosts instead of units of selection during a CRISPR-MACE campaign, selection circuits are highly resistant to cheating or escape. Moreover, beneficial variants rapidly fix to high levels during CRISPR-MACE campaigns, allowing easy nomination of top candidates for confident downstream characterization. This feature of CRISPR-MACE—strong purifying selection—has often proven elusive in other mammalian cell-based directed evolution approaches.
4. The ability to tune selection pressure over the course of evolution campaigns. In this work, fine control was achieved using a small-molecule degradable variant of AcrIIA4. Importantly, CRISPR-MACE’s use of AdProt as a selection marker provides another, generalizable lever for tuning selection pressure. Since every virion must incorporate the AdProt enzyme to be infectious,^50^ there is a stoichiometric need for AdProt. This feature necessarily increases the selection dynamic range more than might otherwise be anticipated for an enzyme-based circuit. Moreover, known AdProt inhibitors^51^ can be employed to further raise selection pressure.
5. The removal of large parts of the adenoviral genome, which avoids the risk of infectious virus recombination.

CRISPR-MACE faced a challenging selection problem in the evolution campaigns reported herein. Cas9 has evolved over billions of years to tightly bind bacteriophage DNA, while AcrIIA4 has evolved over similar timescales to mimic DNA and tightly bind to Cas9. In our implementation of CRISPR-MACE, selected dCas9 variants encoded in the adenoviral genome had to simultaneously avoid binding to their DNA-mimetic inhibitor while still tightly binding to host cell DNA. The resulting interdependent selection pressures allowed us to simultaneously uncover variants of dCas9 with novel DNA binding properties and hitherto inexistent levels of AcrIIA4 resistance. Notably, anti-CRISPRs for diverse Cas9 orthologs are readily available, and some act by diverse mechanisms distinct from inhibiting DNA binding.^52–55^ We predict that implementations of CRISPR-MACE using these orthologous systems will allow the selection of additional novel, complex Cas9 properties. Moreover, there are many opportunities to apply similar MACE-based approaches to optimize the function of other genome targeting agents, including not just miniature CRISPR-Cas proteins, base and prime editors, and epigenetic modifiers, but also other enzyme classes that target the genome.

## METHODS

### Molecular cloning

dCas9–TAD is a fusion of nuclease-inactive *S. pyogenes* Cas9 and the VP64-p65-Rta (VPR) transcriptional activation domain. The gene encoding dCas9–TAD (wild-type or variants) was inserted into the pUC-AdEvolve-DEST plasmid (a custom-made version of AdDest lacking AdProt and AdPol but encoding mCherry; Addgene plasmid #131692)^13^ or a DEST40 plasmid via Gateway cloning (Gateway LR Clonase II Enzyme mix, Thermo Fisher Scientific) with a dCas9–TAD-encoding pENTR1a vector. dCas9-TAD variants were cloned into the pENTR1a vector using the Q5 Site-Directed Mutagenesis Kit (NEB). Derivatives of pUC-AdEvolve-DEST encoding dCas9*–*TAD (dCas9.AdEvolveDest) were used to rescue replication-incompetent adenoviruses as described below, while derivatives of the DEST40 plasmid (dCas9.Dest40) were used for transient transfection-based assays (**Table S5**).

The multi-gRNA expression cassette (U6.gRNA.pLenti in **Table S5**) was assembled via Golden Gate cloning starting with individual gRNA expression plasmids.^40^ The spacers targeted by each of these gRNAs are listed in **Table S6**, as is the Cas9 gRNA scaffold used in this study. The assembled cassette was cloned into a lentiviral transfer plasmid (**Table S5**; to enable lentiviral transduction) via Gibson assembly (NEBuilder HiFi DNA Assembly Master Mix; NEB). The lentiviral transfer plasmid encoding AcrIIA4–csd (*P*_CMV_), as well as plasmids used for stable transfection of AdProt (no promoter) and wild-type AcrIIA4 (*P*_UbC_), were also constructed via Gibson assembly.

For biolayer interferometry (BLI) experiments, dCas9 was obtained from pET28a-Cas9-His (**Table S5**; Addgene # 98158).^56^ In this vector, the wild-type Cas9 gene is placed downstream of a T7 promoter and is followed by a 6× His tag. Introduction of the nuclease-inactivating D10A and H840A substitutions, in addition to the dCas9 variants obtained from directed evolution experiments, was achieved via site-directed mutagenesis using the Q5 Site-Directed Mutagenesis Kit (NEB). AcrIIA4 was cloned into a pET21a vector with a 6× His tag followed by a SUMO tag with a SUMO cleavage site (AcrIIA4.pET21a in **Table S5**).^57^

For luciferase assays, a lentiviral backbone containing CRE-NLuc.PEST (gBlock; IDT) was assembled by Gibson cloning and linearized with BsmBI-v2 for gRNA cloning. The multi-gRNA expression cassette (gRNA.pLenti) was then digested with BsmBI-v2 to release the multi-gRNA sequence, which was ligated into the linearized lentiviral backbone using T4 DNA ligase (the resulting plasmid is referred as pSV23.2 in **Table S5**). Additional constructs, including CRE-NLuc.PEST and EF1a-3×FLAG-AcrIIA4–csd (gBlocks; IDT), were cloned into the lentiviral backbone using Gibson assembly (referred to as pSV23.5 in **Table S5**).

For CASFISH assays, we used a pET302-6His-dCas9-Halo plasmid (**Table S5**; Addgene # 72269)^46^ for bacterial expression of dCas9-HaloTag. dCas9 variants were then generated via site-directed mutagenesis using the Q5 Site-Directed Mutagenesis Kit (NEB).

All final plasmids were verified by full plasmid sequencing (Plasmidsaurus).

### Cell maintenance

With the exception of luciferase assays and lentiviral production, all human cell lines used were derived from the HEK293A cell line (ATCC). Luciferase assays were performed in HEK293T cells (ATCC). Lentivirus was produced in either HEK293T or Lenti-X cells (Takara Bio). Cells were maintained in Dulbecco’s Modified Eagle Medium (Corning) supplemented with 1% v/v penicillin-streptomycin (Corning), 10% fetal bovine serum (Corning), and L-glutamine (Corning). Cells were cultured in either 10-cm or 15-cm dishes and were subcultured at a 1:10 dilution 1–2 times per week. When applicable, the culture medium was supplemented with Hygromycin (5 μg/mL active concentration; Gibco), Puromycin (1 μg/mL active concentration; Gibco), Blasticidin (10 μg/mL active concentration; Gibco), or any combination of the three antibiotics, to select for and maintain transgenes. For the CASFISH assay, mouse embryonic fibroblasts (MEFs; ATCC SCRC-1008) were cultured in (DMEM; Gibco, Cat. No. 10564029) supplemented with 10% (v/v) fetal bovine serum (FBS; Gibco) and 1% v/v penicillin–streptomycin. All cells were regularly assayed for mycoplasma contamination using the MycoStrip Mycoplasma Detection Kit (InvivoGen).

### Cell line generation

We found that the complexity of our circuit, requiring optimized expression of multiple constructs, demanded construction of stable cell lines rather than simple transient transfection of the individual pieces. Thus, to generate custom-engineered cell lines for directed evolution campaigns, lentiviral vectors (for stable transduction) were generated by transfecting a confluent 10-cm plate of Lenti-X cells with plasmids from a third-generation lentiviral packaging system.^58^ Transfection mixtures comprising 1.3 μg of pMDLg/pRRE (Addgene #12251), 2.6 μg of pRSV-Rev (Addgene #12253), 2.6 μg of pMD2.G (Addgene #12259), and 3.3 μg of the transgene-encoding transfer vector (**Table S5**) were mixed with 30 μL of TransIT-Lenti (Mirus Bio) and 1 mL of Opti-MEM before being applied to Lenti-X cells. After 72 h, the culture medium was harvested, centrifuged to remove cell debris, and 3 mL of the supernatant (supplemented with 4 μg/mL Polybrene; MilliporeSigma) was used to transduce a confluent 10-cm dish of the appropriate cells. Selection for stable integrants was initiated 72 h post-transduction with the application of fresh medium containing appropriate selection antibiotics.

Some transgenes in stable cell lines were instead introduced by transfection and subsequent selection. In these cases, cells were transfected with 10 μg of the transgene-encoding plasmids (**Table S5**). 48 h post-transfection, fresh medium containing appropriate selection antibiotics was applied. See **Supplementary Note 1** for a more detailed description of the cell lines used in CRISPR-MACE and the process of their generation.

To generate cell lines for luciferase assays, Lentiviral particles were generated in HEK293T cells by co-transfecting psPAX2 (Addgene #12260), pMD2.G (Addgene #12259), and either pSV23.2 or pSV23.5 at a 2:1:3 ratio, using TransIT®-LT1 reagent (MirusBio) at 5 μg total DNA per million cells. Conditioned medium was collected 48 h post-transfection, filtered through 0.45 μm PES filters (Millipore Sigma), and used to transduce HEK293T cells in the presence of 10 μg/mL polybrene (Millipore Sigma). Transduced cells were selected with 2 μg/mL puromycin for 72 h, followed by maintenance in medium containing 1 μg/mL puromycin for routine culture.

### Adenovirus production

dCas9–TAD-encoding pUC-AdEvolve-DEST plasmids (dCas9.AdEvolveDest; **Table S5**) were PacI-digested and ∼10 μg of the linearized vector was transfected onto confluent 10-cm plates of producer cells. Red fluorescence was used to monitor the extent of viral replication and cell culture media was supplemented every 2–3 d until viral plaques were observed (∼2–3 weeks). Once viral proliferation lysed all cells (approximately one week after the appearance of the first plaques), cells and media were harvested and subjected to three freeze–thaw cycles (−80 °C for at least 30 min, 37 °C for ∼15 min). Cell debris was removed by centrifugation and supernatant was applied to a fresh plate of cells to increase viral titer. This amplification step was repeated until viral proliferation took <1 week on producer cells, usually requiring 2–3 amplification steps.

### RT-qPCR

95% confluent cells (at 12-well scale) were transfected with 1 μg of dCas9.Dest40 (**Table S5**) using TransIT Lenti (Mirus Bio) transfection reagent and Opti-MEM (Gibco) carrier medium. 2 d post-transfection, RNA was extracted using E.Z.N.A total RNA kit I and Homogenizer Mini Columns (Omega Bio-Tek). 1 μg of RNA was used to generate 20 μL of cDNA using the High Capacity cDNA Reverse Transcription Kit (Applied Biosystems). cDNA was diluted with nuclease-free water to 80 μL, and 2 μL of the diluted cDNA was combined with 8 μL of SYBR Green master mix (KAPA/Roche) containing forward and reverse primers for a 10 μL qPCR reaction at a final primer concentration of 125 nM. Each sample (biological replicate) was run in technical triplicate or quadruplicate in a Roche LightCycler 480. AdProt, AcrIIA4, dCas9–TAD, or gRNA transcript levels were normalized to transfection of empty and/or control plasmids. For *in cellula* CRISPRa experiments, AdProt transcript levels were additionally normalized against dCas9–TAD transcript levels. Data were plotted using GraphPad Prism (version 10.4.2). qPCR primers used in this study are listed in **Table S7**.

### CRISPRa luciferase assay

HEK293T cells stably expressing the multi-gRNA cassette (from U6.gRNA.pLenti) targeting AdProt and CRE-NLuc.Pest were nucleofected with a mixture of dCas9.Dest40 (300 ng; **Table S5**) along with AcrIIA4–csd.pLenti (50 ng; **Table S5**) for experiments involving AcrIIA4 inhibition. For each condition, 0.3 million cells were resuspended in 20 µL of SF nucleofection buffer (Lonza) and subjected to nucleofection using the CM-130 program on the Lonza 4D-Nucleofector. Following nucleofection, cells were recovered at rt for 15 min, supplemented with culture medium, and seeded at 30,000 cells/well in 96-well plates. Cells were incubated for 24 h prior to viability and luminescence assays. For normalization, viability was first assessed using PrestoBlue reagent (1:10 dilution in media; Thermo Fisher Scientific, A13262) and fluorescence was read on a Spectramax plate reader. Luminescence was measured following the addition of 10 µM furimazine (Promega) in culture medium, after a 10 min incubation at 37 °C, using an PHERAstar FSX plate reader.

### Adenoviral titering

For plaque assays, producer cells (**Supplementary Note 1**) were plated near confluency on 6-well plates. The following day, 100 μL of various dilutions of virus-containing media were applied to cells. 6 h post-infection, culture media was replaced with media containing 0.4% agarose. Infected cells were monitored to observe the growth of viral plaques. Once plaques were macroscopically visible (∼3 weeks), cells were stained with 3-(4,5-dimethylthiazol-2-yl)-2,5-diphenyltetrazolium bromide (50 mg in 10 mL PBS; Millipore Sigma) and developed for 2 h. Following staining, plaques (which appear as clear holes in a blue monolayer) were counted, but only on wells where the total number of plaques was between 5–50 to ensure sufficient data for quantitation. Titer was calculated as: (number of plaques observed) ÷ (0.1 × dilution factor) and reported in plaque-forming units per milliliter (PFU/mL).

For flow cytometry-based adenovirus titering experiments, wild-type AdPol-expressing cells were plated on 12-well plates. The following day, known volumes of virus-containing media (diluted or not) were applied to cells. 72 h post-infection, cells were washed with media twice, stained with 4′,6-diamidino-2-phenylindole (DAPI; Thermo Fisher Scientific), trypsinized, and transferred to strainer-capped flow cytometry tubes. Samples were analyzed on a BD LSR II analyzer for fluorescent protein (mCherry) expression. Titers were determined as: (fraction of fluorescent cells × total number of cells per well) ÷ (volume of virus-containing media added in mL × dilution factor) and reported in infectious units per milliliter (IU/mL). To minimize inclusion of doubly infected cells, data were only considered when fluorescent cells comprised ≤30% of a given sample.

### Evolution workflow

Evolution campaigns were initiated by seeding polyclonal or monoclonal selector cells at 25% confluency (∼2.5 · 10^6^ cells) on 10-cm dishes the day before infection. Just before transduction, the culture medium was replaced with fresh media containing 10 μM pomalidomide. Selector cells (polyclonal as well as monoclonal) were then transduced with a homogenous population of wild-type dCas9–TAD encoding adenovirus at a multiplicity of infection (MOI) of 10 (campaign #1, polyclonal selector cells) or 14 (campaign #2, monoclonal selector cells). High MOIs and concentrations of pomalidomide were used at the onset of both evolution campaigns to ensure that even wild-type dCas9 encoding virions could replicate in an error-prone fashion (via EpPol). About 2–3 d post-infection, the media on cells was swapped with fresh media containing 10 μM pomalidomide. Upon complete viral proliferation-induced cell death, culture medium containing cell debris and adenovirus was harvested and subjected to three freeze–thaw cycles (−80 °C for at least 30 min, 37 °C for about 15 min). After freeze–thaws, samples were centrifuged at 1000 rpm for 5 min to pellet cellular debris, and virus-containing media was transferred to a new container for storage.

Subsequent passages were initiated by replacing the culture medium on 25% confluent selector cells with fresh culture medium containing pomalidomide (at some researcher-defined concentration, usually lower than used in the previous round) and transferring a small aliquot of viral media from the previous round (note that this media also contained some amount of pomalidomide, presumably the same concentration as in the previous round subject to chemical degradation of the pomalidomide). The final concentration of pomalidomide was estimated by taking pomalidomide in both media fractions (fresh and from the previous round) into account. Cells were supplemented with fresh medium containing that appropriate concentration of pomalidomide every 4–6 d. This viral passaging and pomalidomide passaging procedure was repeated until evolving viral populations were able to replicate robustly even when the concentration of pomalidomide was vanishingly small.

Note that the effective MOI for both evolution campaigns precipitously declines after the earliest passages. At sufficiently high selection pressures, viruses encoding initially prevalent but non-functional dCas9 variants (e.g., wild-type dCas9) are unable to replicate. On the other hand, functional variants, resulting from error-prone replication, initially exist at very low frequencies. Thus, a large “viral die-off” is seen whenever selection pressure is increased (concomitant to a decrease in pomalidomide), leading to a sharp drop to MOI < 1.

### Nanopore sequencing

Adenoviral DNA was isolated from ∼200 μL of virus-containing media using the Quick-DNA/RNA Viral Kit (Zymo Research). The dCas9 open reading frame was then PCR-amplified using Q5 High-Fidelity DNA Polymerase (NEB) and dCas9-specific primers (**Table S7**). The amplicons were bead-purified and prepared for nanopore sequencing using Oxford Nanopore’s Native Barcoding Kit 96 V14 (SQK NBD114.96). Following nanopore sequencing on a MinION10.4.1 flow cell, FASTA/ POD5 formatted files were base-called using a custom Google Colab script invoking Oxford Nanopore’s guppy (ont-guppy_6.5.7) or dorado-0.8.2 base-callers. Fastq files were filtered using a Q20 cutoff to exclude low-quality reads via the ‘NanoFilt’ module (NanoFilt 2.8.0). Data were aligned to the dCas9 reference sequence using Oxford Nanopore’s general-purpose alignment program ‘Minimap2’ (minimap2/2.24). Finally, variants and their respective allele frequencies were identified post-mapping using the ‘LoFreq’ variant caller (2.1.3.1).^59^

### Western blotting

Wild-type AcrIIA4-expressing cells, polyclonal selector cells, monoclonal selector cells, and control cells (the ‘parental’ cell line) were plated on 6-well plates at a density of 5 × 10^6^ cells/well. 3 d post-plating, cells were washed with phosphate-buffered saline and lysed on the plate for 5 min in M-PER (Thermo Fisher Scientific) containing cOmplete™, EDTA-free Protease Inhibitor Cocktail Tablets (Roche). The lysate was vortexed for 30 sec, incubated on ice for 15 min, then clarified by centrifugation at 21,000 × *g* for 20 min at 4 °C. Total protein concentrations were quantified by Bradford assays (Thermo Fisher Scientific). 20 μg of total protein per sample was treated with 6× gel loading buffer (300 mM Tris at pH 6.8, 30% glycerol, 6% SDS, 10% (w/v) bromophenol blue, and 100 mM DTT) and boiled for 10 min. Samples were separated on 4–12% NuPAGE Bis-Tris Mini Protein Gels (Thermo Fisher Scientific), and protein bands were transferred to a nitrocellulose membrane. Immunoblots were probed with an anti-AcrIIA4 primary antibody (Abcam ab273439; 1:1000) and an anti-β-actin primary antibody (Millipore Sigma A1978; 1 μg/mL), followed by an 800 CW anti-mouse secondary antibody (LI-COR 926-32210; 1:5000). Immunoblots were imaged using an Odyssey infrared imager (LI-COR).

### Capillary electrophoretic immunoblotting

HEK293T cells stably expressing 3×FLAG-AcrIIA4–csd (from pSV23.5; **Table S5**) were treated with varying concentrations of pomalidomide for 12 h. For proteasomal inhibition controls, cells were pretreated with MG132 (1 μM) for 2 h prior to pomalidomide exposure. Following treatment, cells were pelleted by centrifugation at 500 × *g* for 5 min and lysed on ice for 30 min in M-PER supplemented with cOmplete™, EDTA-free Protease Inhibitor Cocktail Tablets (Roche). Lysates were clarified by centrifugation at 20,000 × *g* for 20 min at 4 °C, and protein concentration was quantified using the bicinchoninic acid assay (Thermo Fisher). Lysates were diluted to 1 μg/μL in 1× fluorescent master mix (EZ standard pack I), and 3 μL of each sample was loaded per capillary (12–230 kDa cartridge) and analyzed on a Jess Simple Western System (ProteinSimple). AcrIIA4–csd degradation was assessed using an anti-FLAG primary antibody (Cell Signaling Technologies 2368S at 1:50 dilution) with an anti-rabbit HRP secondary antibody (ProteinSimple 042-206; undiluted). Loading was assessed using an anti-tubulin primary antibody (Cell Signaling Technologies, 3873S at 1:50 dilution) with an anti-mouse secondary NIR antibody (ProteinSimple 043-821; 1:20). Luminescence and fluorescence signals were processed using Compass for SW software (v6.1.0).

### Protein purification

For the purification of dCas9, BL21 cells (Thermo Fisher Scientific) harboring pET28a-Cas9-His or pET302-6His-dCas9-Halo plasmids (**Table S5**) were grown o/n in TB media at 37 °C and then used to inoculate a secondary 1 L culture. These cultures were grown at 37 °C until they reached OD_600_ = 0.6, at which point 0.3 mM isopropyl-β-D-1-thiogalactosidease (IPTG; Thermo Fisher Scientific) was added at an induction temperature of 18 °C. Following an additional 16 h growth, cultures were centrifuged at 6000 rpm for 10 min at 4 °C in a Sorvall LYNX 6000 to harvest cell pellets, which were subsequently resuspended in buffer (25 mM Tris-Cl at pH 7.4, 1 M KCl, 5 mM MgCl_2_, 1 mM TCEP, and 20% glycerol). All subsequent steps were performed at 4°C.

Cell lysis was initiated by adding 1 mg/ml of lysozyme, 0.1 mg/ml of DNase I (Stem Cell Technologies), 0.5 mM PMSF (Thermo Fisher Scientific), and 1× protease inhibitor cocktail (Pierce; one tablet/mL lysis buffer) to the resuspended pellets. This mixture was incubated for 1 h before adding 0.5% Triton X-100 (Millipore Sigma) and incubating for an additional 15 min. Ultrasonic lysis was carried out using a 250-watt processor (Fisher Scientific) with a 6-mm diameter probe and applying a cycle of 10 a ON and 15 a OFF for 15 min. After centrifuging the lysate to remove cell debris, the cleared lysate was subjected to affinity chromatography by incubating it with 1.5 mL of pre-equilibrated His-Pur Cobalt nitriloacetic acid (Co-NTA) resin (Thermo Fisher Scientific) for 2 h before washing 3× with 15 column volumes of washing buffer (25 mM Tris-Cl at pH 7.4, 100 mM KCl, 5 mM MgCl_2_, 1 mM TCEP, 40 mM imidazole, and 20% glycerol) and eluting with elution buffer (25 mM Tris-Cl at pH 7.4, 100 mM KCl, 5 mM MgCl_2_, 1 mM TCEP, 300 mM imidazole, and 20% glycerol).

The eluted dCas9 protein was concentrated using 100 kDa cut-off Amicon spin concentrators (Millipore Sigma), centrifuged at 3450 rpm for 15 min, and subjected to gel filtration chromatography using an AKTA Pure (Cytiva). A final purification step was undertaken using a HiLoad Superdex-200 size exclusion chromatography column (Cytiva) at a flow rate of 1 mL/min using running buffer (25 mM Tris-Cl at pH 7.4, 100 mM KCl, 5 mM MgCl_2_, 1 mM TCEP, and 20% glycerol). Protein-containing fractions were assessed using NuPAGE Bis-Tris gels with a 4–12% gradient (Thermo Fisher Scientific) to determine their final purity. Purified dCas9 samples were sterile-filtered through a 0.22 µm PVDF membrane filter (Millipore Sigma), quantified via BCA assay (Pierce), snap-frozen in N_2_(l), and stored in aliquots at −80 °C.

For the purification of AcrIIA4, BL21 cells harboring AcrIIA4.pET21a (**Table S5**) were grown o/n in TB media at 37 °C and then used to inoculate a secondary 1 L culture. Once the secondary culture reached OD_600_ = 0.6, 0.5 mM IPTG was added at 18 °C. Following an additional 16 h growth, the AcrIIA4-expressing culture was lysed and affinity purified via a similar protocol to that described above for dCas9 purification. However, the purified His6×-SUMO-AcrIIA4 polypeptide was additionally subjected to a tag cleaving reaction by incubating it with SUMO protease and Ulp-1 (Thermo Fisher Scientific) in a 10:1 ratio in tag cleaving buffer (25 mM Tris-Cl at pH 7.4, 100 mM KCl, 1mM TCEP) o/n at 4 °C with gentle rocking. The following day, this mixture was incubated with 0.5 mL of pre-equilibrated His-Pur Cobalt nitriloacetic acid (Co-NTA) resin for 30 min and tag-cleaved AcrIIA4 was collected as the unbound fraction.

Tag-free AcrIIA4 was concentrated using 3 kDa cut-off Amicon spin concentrators (Millipore Sigma), centrifuged at 3450 rpm for 15 min, and subjected to gel filtration chromatography using an AKTA pure system (Cytiva). A final purification step was undertaken using a HiLoad Superdex-75 size exclusion chromatography column (Cytiva) at a flow rate of 0.8 mL/min using running buffer (25 mM Tris-Cl at pH 7.4, 100 mM KCl, 5 mM MgCl_2_, 1 mM TCEP, and 20% glycerol). Protein-containing fractions were assessed using NuPAGE Bis-Tris gels with a 4–12% gradient (Thermo Fisher Scientific) to determine their final purity. Purified AcrIIA4 samples were sterile-filtered through a 0.22 µm PVDF membrane filter, quantified via BCA assay, snap-frozen in N_2_(l), and stored in aliquots at −80 °C.

### Biolayer interferometry (BLI) assays

Ribonucleoprotein (RNP) complexes were prepared by incubating 25 nM of each respective dCas9 variant (prepared as described above) with 30 nM of gRNA (preparation described below) in 1× assay buffer (20 mM Tris-Cl sy pH 7.4, 100 mM KCl, 5 mM MgCl_2_, 1 mM TCEP, 0.01% Tween-20, 50 µg/ml heparin) at 4 °C for 15 min. Control complexes were prepared similarly by substituting gRNA-containing buffer for 1× assay buffer.

The gRNA component of the RNP complexes was prepared by amplifying linear DNA fragments containing the T7 RNA polymerase promoter sequence upstream of the desired gRNA spacer and backbone sequences (**Table S6**) using Q5 Hot Start MasterMix (NEB) and appropriate forward and reverse primers (**Table S7**). These fragments were then concentrated using Mini Elute columns (Qiagen) and *in vitro* transcribed using the HiScribe T7 High Yield RNA Synthesis Kit (NEB) at 37 °C for 14–16 h with 200 ng of linear template per 20 μL reaction. Transcribed gRNA was purified using the MEGAClear Transcription Clean-Up Kit (Thermo Fisher Scientific), according to the manufacturer’s instructions. The concentration of the purified gRNA was determined by diluting an aliquot 1:100 in TE buffer (10 mM Tris-HCl at pH 8; 1 mM EDTA) and measuring absorbance at 260 nm. The RNA concentration in µg/mL was calculated using the following formula: Abs_260nm_ × dilution factor × 40 μg/mL. Quantified gRNA was stored in aliquots at −80 °C.

The biotinylated dCas9 dsDNA substrate was prepared by annealing 10 μM forward and reverse oligos (**Table S7**) in a mixture of annealing buffer (20 mM Tris-Cl at pH 7.5, 100mM KCl, 5 mM MgCl_2_) and nuclease-free water in a PCR tube. This mixture was heated to 95 °C for 5 min, then cooled to 25 °C at 0.1 °C/s. The resulting dsDNA substrate, which had a concentration of 10 μM, was then diluted to 1 μM in 1× assay buffer and stored at −20 °C.

All AcrIIA4 binding assays were performed in 1× assay buffer (20 mM Tris-Cl at pH 7.4, 100 mm KCl, 5 mM MgCl_2_, 1 mM TCEP, 0.01% Tween-20, 50 µg/ml heparin) using an 8-channel Octet R8 (Sartorius) at 25 °C. 300 nM of biotinylated substrate DNA was immobilized onto streptavidin (SAX) biosensors (Sartorius) for 300 s up to a response of 0.8-1 nm. The reference biosensors were immobilized similarly, except they were dipped in 1× assay buffer instead of substrate DNA. All biosensors were blocked with 100 µM biocytin (Millipore Sigma) supplemented in 1× assay buffer for 300 s and a stable baseline was achieved for another 120 s. The respective RNP complexes were titrated with a six-point dose of AcrIIA4 from 1–10 nM and association and dissociation kinetics were measured for 1200 s each in 1× assay buffer. All kinetics experiments were performed in biological duplicate.

All DNA binding experiments were performed in 1× assay buffer (20 mM Tris-Cl at pH 7.4, 100 mm KCl, 5 mM MgCl_2_, 1 mM TCEP, 0.01% Tween-20, 50 µg/ml heparin) using an 8-channel Octet R8 (Sartorius) at 25 °C. 150 nM of biotinylated substrate DNA was immobilized onto SAX biosensors (Sartorius) for 300 s up to a response of 0.5–0.6 nm. Control biosensors (**Figures S8b, d, f, h, j, l and S9b, d, f, h, j**) were prepared in a similar manner; except they were dipped in 1× assay buffer instead of substrate DNA.

For the DNA binding experiments, all biosensors were blocked with 100 µM biocytin (Millipore Sigma) supplemented in 1× assay buffer for 300 s, and a stable baseline was achieved for another 120 s. The RNP complexes prepared with dCas9 (wild-type or variants) were two-fold serially 6×, starting from 100 nM (seven concentrations tested). Binding (association and dissociation) kinetics were measured for 1200 s each, with 180 s baseline in 1× assay buffer. Control kinetics experiments were performed in a similar way using control biosensors. All kinetics experiments were performed in biological duplicate.

Raw data were processed using the inbuilt ‘Octet BLI analysis 12.2 software from Sartorius. The reference biosensors were subtracted using double referencing, and subtracted data were aligned with baseline for each experiment. Data were further aligned using inter-step correction and processed for Savitzky-Golay filtering. Kinetic parameters such as *k*_on_, *k*_off_, and *K*_d_ were determined by globally fitting the association and dissociation phases from the DNA binding assays (no AcrIIA4) to a 1:1 binding model. To calculate IC_50_ values (from AcrIIA4 binding experiment), all binding curves for each experiment were normalized against the “RNP, no AcrIIA4” condition (set as 100% binding) and plotted as a function of AcrIIA4 concentration. The resulting dose-response curves were fitted using non-linear regression to derive IC_50_ values. The final data were plotted using GraphPad Prism (version 10.2.3). Each kinetics experiment included full biological replicates from independent dCas9 preparations (*n* = 2), and the IC_50_ values are reported as means with standard deviation.

### CASFISH Assay

CASFISH experiments were performed as previously described with modifications.^46^ The methods for dCas9–HaloTag (from pET302-6His-dCas9-Halo; **Table S5**) protein expression and purification, as well as sgRNA preparation (**Tables S6** and **S7**), are described in the BLI experimental section. dCas9–HaloTag and variant proteins were diluted to 2 µM in 50 mM HEPES at pH 7.5, 150 mM KCl, and 1 mM TCEP, and incubated with an eight-fold molar excess of Janelia Fluor 646 (JF646)–Halo ligand conjugate (Tocris) for 30 min at rt, followed by o/n incubation at 4 °C. Unreacted JF646–Halo ligand was removed using 40 kDa molecular weight cut-off Zeba spin desalting columns (Thermo Scientific). Labeled proteins were eluted in 50 mM HEPES at pH 7.5, 150 mM KCl, 1 mM TCEP, and 10% (v/v) glycerol.

Mouse embryonic fibroblasts were seeded at passage 2 into black 96-well PhenoPlates (Revvity) at a density of 15,000 cells/well. After o/n adherence, cells were fixed with a pre-chilled 1:1 mixture of MeOH and acetic acid for 20 min at −20 °C. Fixed cells were washed 3× with PBS (5 min each, with gentle shaking) and incubated with blocking buffer (20 mM HEPES at pH 7.5, 100 mM KCl, 5 mM MgCl₂, 5% (v/v) glycerol, 1% BSA, 0.1% Tween-20, and freshly prepared 1 mM DTT) for 15 min at 37 °C. CASFISH ribonucleoprotein (RNP) probes were assembled by mixing fluorescently labeled dCas9–HaloTag (100 nM) with sgRNA (400 nM) in blocking buffer for 10 min at rt, followed by incubation on ice. The assembled RNP complexes (1 nM) were added to the cells and incubated for 10 min at 37 °C. Cells were subsequently washed 3× with blocking buffer. Prior to imaging, samples were stained with Hoechst 33342 (0.5 µg/mL) for 5 min.

High-throughput fluorescent imaging of JF646 (Alexa 647 channel) and Hoechst 33342 was performed using a 60× water objective on an automated Opera Phenix High-Content Imaging System (PerkinElmer), and image analysis was conducted with Harmony software (v4.9, PerkinElmer) across a z-stack comprising 0.5 μm sections. Built-in algorithms were used to quantify the number and intensity of nuclear spots in the Alexa 647 channel for each detected nucleus in the maximum projection image of the z-stack. Mean spot intensity per cell was calculated as the average signal intensity of all spots within individual nuclei. Data were plotted as distributions of nuclear intensities from at least 200 cells across 30 fields of view per condition using GraphPad Prism (version 10.4.2).

## DATA, MATERIALS, AND SOFTWARE AVAILABILITY

Sequencing data are available at NCBI Bioproject ID PRJNA1379435. Other study data are included in the article and/or supporting information. Plasmids are listed in **Table S5** and the corresponding sequences for generated plasmids will be deposited to GenBank upon article acceptance.

## ACKNOWLEDGEMENTS

This work was supported by the NIH (R35GM136354 to M.D.S. and R01GM132825, R01DK132900, and R01GM137606 to A.C.), the National Science Foundation Division of Chemistry Molecular Foundations of Biotechnology Research Grant 2330699 (to M.D.S.), the Broad Institute’s Merkin Institute of Transformative Technologies in Healthcare Grant (to A.C. and M.D.S.), MIT Deshpande Center for Technological Ignition and Innovation Grants (to M.D.S.), and DARPA (N66001-17-2-4055 to A.C.). A.A.M. was supported by a Future of Science Fellowship from the MIT School of Science, a Robert J. Silbey Graduate Fellowship from the MIT Department of Chemistry, and a BroadIgnite Fellowship. A.L.S., A.M.B., S.J.H., and C.M.L. were supported by National Science Foundation Graduate Research Fellowships. K.K. was supported by a Natural Sciences and Engineering Research Council of Canada Post-Doctoral Fellowship (587836-2024). C.L. was supported by the Damon Runyon Cancer Research Foundation as a Suzanne and Bob Wright Fellow (DRG-2539-24). This work was also supported in part by the National Cancer Institute under award P30-CA14051 and the National Institute of Environmental Health Sciences under award P30-ES002109.

