## Supporting Information for "Anti-CRISPR-mediated continuous directed evolution of CRISPR-Cas9 in human cells"


| PAGE | CONTENTS |
|---|---|
| **S1** | Table of Contents |
| **S2** | Supplementary Notes |
| **S3–12** | Supplementary Tables |
| **S13–26** | Supplementary Figures |
| **S27** | Supplementary References |

### SUPPLEMENTARY NOTES

**Supplementary Note 1.** Description of cell lineages for the generation of selection cell lines and other cell lines used in CRISPR-MACE.

The 'producer' cell line was generated via lentiviral transduction of HEK293A cells with AdPol (expressed via $P_{CMV}$) and two AdProt cassettes (one expressed via $P_{CMV}$ and the second expressed via $P_{TetO}$). This cell line constitutively trans-complements all the essential missing adenoviral genes (AdProt, AdPol, and the E1 region, but not the E3 region that is only required for immune escape *in vivo*) and is used to rescue or propagate replication-incompetent virus in the absence of selection pressure.[1] Additional production of AdProt can be driven by the inducible AdProt cassette, but activation of that cassette was not required for the studies reported here. These cells are also used for plaque assays to determine adenoviral titer.

Selector cell lines used as hosts during virus-assisted evolution were generated via a multistep process beginning with the lentiviral transduction of HEK293A cells with an error-prone variant of AdPol (EpPol) whose expression is driven by $P_{CMV}$. Integration of EpPol was followed by the stable transfection of an AdProt-encoding plasmid with only a minimal CMV promoter upstream of AdProt (AdProt.pGL4; **Table S5**). After establishing a polyclonal population through the addition of Hygromycin (5 µg/mL active concentration), monoclonal lines were generated by dilution and manually picking monoclonal islands. The monoclonal cell line that had the lowest basal expression of AdProt was selected as a 'parental' cell line into which other elements were subsequently integrated. An AcrIIA4–csd cassette whose expression was driven by $P_{CMV}$ (AcrIIA4–csd.pLenti; **Table S5**) was one such element.[2] Integration of AcrIIA4–csd was performed via lentiviral transduction of AcrIIA4–csd.pLenti. Once an AcrIIA4–csd-encoding polyclonal population was established through the addition of Puromycin (1 µg/mL active concentration), monoclonal lines were generated and the monoclonal cell line with the highest expression of AcrIIA4–csd (measured by reverse transcription quantitative PCR; RT-qPCR) was lentivirally transduced with the multi-gRNA cassette (U6.gRNA.pLenti; **Table S5**) and a polyclonal population was established through the addition of Blasticidin (10 µg/mL active concentration). The resulting polyclonal cell line as well as a monoclonal cell line derived from it that displayed the highest expression of most gRNAs were dubbed the polyclonal and monoclonal selector cells, respectively.

The wild-type AcrIIA4-expressing cell line (used to assess resistance to AcrIIA4 *in cellula* via RT-qPCR) was similarly generated via lentiviral transduction of the multi-gRNA cassette (U6.gRNA.pLenti; **Table S5**) into the EpPol- and AdProt-encoding 'parental' cell line. However, wild-type AcrIIA4 was then stably transfected (cistron placed under the control of $P_{UbC}$; UbC.WT-AcrIIA4.pLenti in **Table S5**) to ensure this cell line's AcrIIA4 expression levels matched that of AcrIIA4–csd in the selector cell lines.

### SUPPLEMENTARY TABLES

**Table S1.** List of nucleotide and amino acid substitutions observed in the dCas9 locus in Campaign #1 (conducted in the polyclonal selector cells).

| Passage | Reference codon | Nucleotide change | Variant codon | Non-reference allele frequency | Amino acid change |
|---|---|---|---|---|---|
| 5 | GGC | G→A | G**A**C | 0.056522 | Gly→Asp (G12D) |
| 6 | GGC | G→A | G**A**C | 0.899381 | Gly→Asp (G12D) |
| | GCC | G→A | **A**CC | 0.075443 | Ala→Thr (A208T) |
| 7 | GGC | G→A | G**A**C | 0.987378 | Gly→Asp (G12D) |
| | GCC | G→A | **A**CC | 0.061899 | Ala→Thr (A208T) |
| 8 | GGC | G→A | G**A**C | 0.984415 | Gly→Asp (G12D) |
| 9 | GGC | G→A | G**A**C | 0.98607 | Gly→Asp (G12D) |
| 10 | GGC | G→A | G**A**C | 0.988018 | Gly→Asp (G12D) |
| | CGC | C→T | **T**GC | 0.082037 | Arg→Cys (R400C) |
| | AAG | G→A | AA**A** | 0.074876 | Lys→Lys (K506K) |
| | TGT | T→A | **A**GT | 0.072853 | Cys→Ser (C574S) |
| | GCC | C→T | G**T**C | 0.196416 | Ala→Val (A764V) |
| | GTG | G→A | GT**A** | 0.051967 | Val→Val (V1146V) |
| 11 | GGC | G→A | G**A**C | 0.988445 | Gly→Asp (G12D) |
| | GCC | C→T | G**T**C | 0.213623 | Ala→Val (A764V) |
| 12 | GGC | G→A | G**A**C | 0.986546 | Gly→Asp (G12D) |
| | ATT | A→T | **T**TT | 0.116157 | Ile→Phe (I300F) |
| | CGC | C→T | **T**GC | 0.232227 | Arg→Cys (R400C) |
| | CAC | A→G | C**G**C | 0.137987 | His→Arg (H420R) |
| | AAG | G→A | AA**A** | 0.227456 | Lys→Lys (K506K) |
| | GTG | G→A | **A**TG | 0.112959 | Val→Met (V561M) |
| | GCC | C→T | G**T**C | 0.734435 | Ala→Val (A764V) |
| | GTG | G→A | GT**A** | 0.117725 | Val→Val (V1146V) |

**Table S1 continued:**

| Passage | Reference codon | Nucleotide change | Variant codon | Non-reference allele frequency | Amino acid change |
|---|---|---|---|---|---|
| 13 | GGC | G→A | G**A**C | 0.990502 | Gly→Asp (G12D) |
| | GGC | C→T | GG**T** | 0.11966 | Gly→Gly (G283G) |
| | ATT | A→T | **T**TT | 0.033475 | Ile→Phe (I300F) |
| | CGC | C→T | **T**GC | 0.097342 | Arg→Cys (R400C) |
| | CAC | A→G | C**G**C | 0.082721 | His→Arg (H420R) |
| | GAT | A→G | G**G**T | 0.05756 | Asp→Gly (D428G) |
| | AAG | G→A | AA**A** | 0.095677 | Lys→Lys (K506K) |
| | GTG | G→A | **A**TG | 0.034539 | Val→Met (V561M) |
| | GCC | C→T | G**T**C | 0.936463 | Ala→Val (A764V) |
| | GGA | G→A | G**A**A | 0.031732 | Gly→Glu (G773E) |
| | TCT | T→A | **A**CT | 0.079864 | Ser→Thr (S845T) |
| | AGG | G→A | A**A**G | 0.162712 | Arg→Lys (R919K) |
| | GTT | G→A | **A**TT | 0.042894 | Val→Ile (V955I) |
| | GTC | G→C | **C**TC | 0.029184 | Val→Leu (V1139L) |
| | GTG | G→A | GT**A** | 0.050597 | Val→Val (V1146V) |
| | AGC | C→T | AG**T** | 0.025072 | Ser→Ser (S1159S) |
| | AGC | G→A | AA**C** | 0.010651 | Ser→Asn(S1240N) |
| | CAC | C→T | CA**T** | 0.009269 | His→His (H1311H) |
| 14 | GGC | G→A | G**A**C | 0.985201 | Gly→Asp (G12D) |
| | GGC | C→T | GG**T** | 0.105108 | Gly→Gly (G283G) |
| | GCC | C→T | G**T**C | 0.961095 | Ala→Val (A764V) |
| | TCT | T→A | **A**CT | 0.076961 | Ser→Thr (S845T) |
| | AGG | G→A | A**A**G | 0.378662 | Arg→Lys (R919K) |
| | GTT | G→A | **A**TT | 0.13873 | Val→Ile (V955I) |
| 15 | GGC | G→A | G**A**C | 0.984459 | Gly→Asp (G12D) |
| | GGC | C→T | GG**T** | 0.089235 | Gly→Gly (G283G) |
| | GCC | C→T | G**T**C | 0.966802 | Ala→Val (A764V) |
| | AGG | G→A | A**A**G | 0.49488 | Arg→Lys (R919K) |
| | GTT | G→A | **A**TT | 0.203037 | Val→Ile (V955I) |
| 16 | GGC | G→A | G**A**C | 0.99309 | Gly→Asp (G12D) |
| | GCC | C→T | G**T**C | 0.977421 | Ala→Val (A764V) |
| | AGG | G→A | A**A**G | 0.594967 | Arg→Lys (R919K) |
| | GTT | G→A | **A**TT | 0.315314 | Val→Ile (V955I) |

**Table S2.** List of nucleotide and amino acid substitutions observed in the dCas9 locus in Campaign #2 (conducted in the monoclonal selector cells).

| Passage | Reference codon | Nucleotide change | Variant codon | Non-reference allele frequency | Amino acid change |
|---|---|---|---|---|---|
| 5 | GGC | G→A | G**A**C | 0.669844 | Gly→Asp (G12D) |
| | CAG | G→T | CA**T** | 0.536981 | Gln→His (Q1221H) |
| 6 | GGC | G→A | G**A**C | 0.62417 | Gly→Asp (G12D) |
| | CAG | G→T | CA**T** | 0.476092 | Gln→His (Q1221H) |
| 7 | GGC | G→A | G**A**C | 0.607508 | Gly→Asp (G12D) |
| | CAG | G→T | CA**T** | 0.447099 | Gln→His (Q1221H) |
| 8 | GGC | G→A | G**A**C | 0.830718 | Gly→Asp (G12D) |
| | CAG | G→T | CA**T** | 0.75118 | Gln→His (Q1221H) |
| 9 | GGC | G→A | G**A**C | 0.934743 | Gly→Asp (G12D) |
| | GCC | G→A | **A**CC | 0.069736 | Ala→Thr (A764T) |
| | CAG | G→T | CA**T** | 0.851877 | Gln→His (Q1221H) |
| 10 | GGC | G→A | G**A**C | 0.981248 | Gly→Asp (G12D) |
| | GCC | G→A | **A**CC | 0.182164 | Ala→Thr (A764T) |
| | CAG | G→T | CA**T** | 0.957479 | Gln→His (Q1221H) |
| 11 | GGC | G→A | G**A**C | 0.98672 | Gly→Asp (G12D) |
| | GCC | G→A | **A**CC | 0.144203 | Ala→Thr (A764T) |
| | CAG | G→T | CA**T** | 0.958225 | Gln→His (Q1221H) |
| 12 | GGC | G→A | G**A**C | 0.986343 | Gly→Asp (G12D) |
| | CGC | C→T | CG**T** | 0.484937 | Arg→Arg (R654R) |
| | GCC | G→A | **A**CC | 0.159317 | Ala→Thr (A764T) |
| | GCC | C→T | G**T**C | 0.562402 | Ala→Val (A764V) |
| | CAG | G→T | CA**T** | 0.986412 | Gln→His (Q1221H) |
| 13 | GGC | G→A | G**A**C | 0.988603 | Gly→Asp (G12D) |
| | CGC | C→T | CG**T** | 0.453961 | Arg→Arg (R654R) |
| | GCC | G→A | **A**CC | 0.155187 | Ala→Thr (A764T) |
| | GCC | C→T | G**T**C | 0.556791 | Ala→Val (A764V) |
| | CAG | G→T | CA**T** | 0.985245 | Gln→His (Q1221H) |
| 14 | GGC | G→A | G**A**C | 0.989096 | Gly→Asp (G12D) |
| | CGC | C→T | CG**T** | 0.381098 | Arg→Arg (R654R) |
| | GCC | G→A | **A**CC | 0.163019 | Ala→Thr (A764T) |
| | GCC | C→T | G**T**C | 0.516765 | Ala→Val (A764V) |
| | CAG | G→T | CA**T** | 0.97453 | Gln→His (Q1221H) |

**Table S3.** AcrIIA4-mediated inhibition ($IC_{50}$ values) of dCas9 variants binding to dsDNA measured using biolayer interferometry experiments. WT = wild-type dCas9, F.C. = fold-change relative to wild-type dCas9. Uncertainty is expressed as standard deviation.

| dCas9 variant | $IC_{50}$ (nM) | F.C. ($IC_{50}$) |
|---|---|---|
| WT | 9.3 ± 6.0 | 1 |
| G12D | 19.6 ± 6.1 | 2.1 |
| A764V | 4.7 ± 0.1 | 0.5 |
| A764T | 6.1 ± 1.5 | 0.7 |
| R919K | 1176.8 ± 266.2 | 126.1 |
| V955I | 456.5 ± 195.7 | 48.9 |
| Q1221H | 6.8 ± 0.1 | 0.7 |
| G12D/A764V (DV) | 225.6 ± 64.6 | 24.2 |
| G12D/Q1221H (DH) | 782.7 ± 165.8 | 83.9 |
| G12D/A764V/Q1221H (DVH) | 940.0 ± 240.5 | 100.7 |
| G12D/A764T/Q1221H (DTH) | 707.9 ± 209.7 | 75.9 |
| G12D/A764V/R919K/V9955I (DVKI) | 2202.5 ± 1174.5 | 236.1 |
| G12D/A764V/R919K/V9955I/Q1221H (DVKIH) | 8308.0 ± 1415.6 | 890.5 |

**Table S4.** List of kinetic parameters for dCas9 variants in DNA binding assay obtained through biolayer interferometry experiments. WT = wild-type dCas9, F.C. = fold-change relative to wild-type dCas9. DV = G12D/A764V; DH = G12D/Q1221H; DVH = G12D/A764V/Q1221H; DVKI = G12D/A764V/R919K/V955I; DVKIH = G12D/A764V/R919K/V955I/Q1221H. Uncertainty is expressed as standard deviation.

| dCas9 variant | $k_{on}$ (× $10^4$), $M^{-1}s^{-1}$ | $k_{off}$ (× $10^{-5}$), $s^{-1}$ | $K_d$ (× $10^{-9}$), M | Residence Time (t), h | F.C. ($K_d$) | F.C. (t) |
|---|---|---|---|---|---|---|
| WT | 0.8 ± 0.0 | 5.4 ± 0.4 | 7.1 ± 0.7 | 5.2 ± 0.4 | 1 | 1 |
| G12D | 1.3 ± 0.1 | 3.0 ± 0.1 | 2.3 ± 0.0 | 9.3 ± 0.5 | 0.3 | 1.8 |
| A764V | 0.5 ± 0.5 | 3.3 ± 0.4 | 9.3 ± 7.3 | 8.5 ± 1.1 | 1.3 | 1.6 |
| R919K | 0.4 ± 0.2 | 8.3 ± 0.4 | 29.4 ± 18.8 | 3.3 ± 0.2 | 4.1 | 0.6 |
| V955I | 0.7 ± 0.0 | 9.8 ± 1.2 | 14.1 ± 0.7 | 2.8 ± 0.4 | 2.0 | 0.6 |
| Q1221H | 1.2 ± 0.1 | 9.6 ± 0.2 | 7.9 ± 1.1 | 2.9 ± 0.1 | 1.1 | 0.6 |
| DV | 0.5 ± 0.1 | 7.2 ± 1.1 | 16.2 ± 4.9 | 3.9 ± 0.6 | 2.3 | 0.8 |
| DH | 1.8 ± 0.0 | 0.7 ± 0.0 | 0.4 ± 0.0 | 39.4 ± 2.1 | 0.1 | 7.6 |
| DVH | 0.9 ± 0.0 | 1.8 ± 0.3 | 2.0 ± 0.2 | 15.4 ± 2.3 | 0.3 | 3.0 |
| DVKI | 0.8 ± 0.1 | 4.3 ± 0.3 | 5.7 ± 0.9 | 6.4 ± 0.4 | 0.8 | 1.2 |
| DVKIH | 1.7 ± 0.2 | 5.0 ± 0.2 | 3.0 ± 0.2 | 5.5 ± 0.2 | 0.4 | 1.1 |

**Table S5.** Key plasmids used in this study are listed below. The corresponding sequences for generated plasmids will be deposited to GenBank upon article acceptance. For gifted plasmids, Addgene and/or literature references are listed.

| Plasmid | Notable component(s) | Description |
|---|---|---|
| AdProt.pGL4 | AdProt, mRNA degradation tag, minimal CMV promoter | Plasmid used to make stable cell line that minimally expressed AdProt in the absence of gRNAs that enable CRISPRa. The AdProt cistron lies downstream of the adenoviral tripartite leader sequence. |
| AcrIIA4–csd.pLenti | AcrIIA4–csd, CMV promoter | Lentiviral transfer plasmid used to make stable cell line that expresses AcrIIA4–csd (AcrIIA4 C-terminally fused to a protein degradation tag). |
| UbC.WT-AcrIIA4.pLenti | WT AcrIIA4, UbC promoter | Lentiviral transfer plasmid used to make stable cell line that expresses wild-type AcrIIA4. This plasmid was stably transfected to make said cell line. |
| U6.gRNA.pLenti | gRNAs (7), U6 promoter | Lentiviral transfer plasmid used to make stable cell lines that express seven gRNAs, all targeting the region right upstream of AdProt (in the pGL4 AdProt plasmid). |
| dCas9.Dest40 | dCas9, TAD (VPR), CMV promoter | Mammalian expression vector encoding dCas9 (wild-type or variants) fused to a transcription activation domain and used to conduct *in cellula* transcription assays. |
| dCas9.AdEvolveDest | dCas9, TAD (VPR), mCherry, modified type 5 adenoviral genome | Engineered adenoviral genome (type 5) encoding dCas9 (wild-type or variants) fused to a transcription activation domain. Genome lacks essential elements for viral replication, including AdProt, AdPol, the E1 region, and the E3 region. It also encodes a fluroscent protein (mCherry) for viral tracking. Transfection of the linearized vector into 'producer' cell lines generates infectious virions. |
| pET28a-Cas9-His Addgene # 98158 (Ref.[3]) | dCas9 (His-tagged), T7 promoter, Lac promoter | Bacterial expression vector encoding dCas9 (wild-type or variants) fused to a His tag (for purification). Vector was used to express and purify dCas9 variants for BLI experiments. |

**Table S5 continued:**

| | | |
|---|---|---|
| AcrIIA4.pET21a (Ref.[4]) | AcrIIA4 (His- and SUMO-tagged), Lac promoter | Bacterial expression vector encoding AcrIIA4 fused to a His tag and a SUMO tag (for purification). Vector was used to express and purify AcrIIA4 for BLI experiments. |
| pSV23.2 | Luciferase-targeting gRNAs (7), Luciferase (promoterless) | Lentiviral transfer vector used to make a stable cell line to assess (1) the *in cellula* dosability of transcriptional activation activity in response to pomalidomide-dependent degradation of AcrIIA4–csd and (2) the *in cellula* transcriptional activation activity of AcrIIA4-resistant dCas9–TAD variants in the absence of AcrIIA4 |
| pSV23.5 | AcrIIA4–csd (3×FLAG-tagged), EF1a promoter, Luciferase (promoterless) | Lentiviral transfer vector used to make a stable cell line to assess (1) the *in cellula* dosability of pomalidomide-dependent AcrIIA4–csd degradation and (2) the *in cellula* transcriptional activation activity of AcrIIA4-resistant dCas9–TAD variants in the presence of AcrIIA4 |
| pET302-6His-dCas9-Halo Addgene # 72269 (Ref.[5]) | dCas9-HaloTag (His-tagged), T7 promoter, Lac promoter | Bacterial expression vector encoding dCas9-HaloTag (wild-type or variants) fused to a His tag (for purification). Vector was used to express and purify dCas9 variants for CASFISH experiments. |

**Table S6.** gRNA spacer and scaffold sequences.

| Component | Sequence (5′→3′) | Notes |
|---|---|---|
| dCas9 gRNA scaffold | GTTTTAGAGCTAGAAATAGCAAGTTAAAA-TAAGGCTAGTCCGTTATCAACTT-GAAAAAGTGGCACCGAGTCGGTGC | Universal scaffold sequence used for all gRNAs |
| AdProt gRNA #1 | GACACCCCATTGACGTCAAT | Spacer sequence located >200 bp upstream of AdProt transcription start site. |
| AdProt gRNA #2 | ACGTCAGCTGCCAGATCCCA | Spacer sequence located >200 bp upstream of AdProt transcription start site. |
| AdProt gRNA #3 | TGGGAGAACAGATCTGGCCT | Spacer sequence located >200 bp upstream of AdProt transcription start site. |
| AdProt gRNA #4 | GCGCTAGCGAGCTCAGGTAC | Spacer sequence located >200 bp upstream of AdProt transcription start site. |
| AdProt gRNA #5 | GAGAACAGATCTGGCCTCGG | Spacer sequence located >200 bp upstream of AdProt transcription start site. |
| AdProt gRNA #6 | TGAAAGACGTCACAGTATGA | Spacer sequence located >200 bp upstream of AdProt transcription start site. |
| AdProt gRNA #7 | AGTATGACGGCCATGGGATC | Spacer sequence located >200 bp upstream of AdProt transcription start site. |
| BLI sgRNA | GACGCATAAAGATGAGACGC | Spacer sequence used to recruit dCas9 to dsDNA substrate during BLI experiments. |
| Pericentromere (Major satellite) sgRNA | CCATATTCCACGTCCTACAG | Spacer sequence used to recruit dCas9 to the centromere of the chromosomes. |

**Table S7.** Oligonucleotides used for RT-qPCR, PCR amplification, or annealed together to make the dsDNA substrate for BLI.

| Number | Sequence (5′→3′) | Notes |
|---|---|---|
| 1 | CGTCGCCTCCTACCTGCT | Forward RT-qPCR primer for RPLP2, a housekeeping gene used as internal control during RT-qPCR. |
| 2 | CCATTCAGCTCACTGATAACCTT | Reverse RT-qPCR primer for RPLP2, a housekeeping gene used as internal control during RT-qPCR. |
| 3 | AAGGCGTCTAACCAGTCACA | Forward RT-qPCR primer used to assess relative AdProt expression. |
| 4 | CGTATTGACTATGGCGCAGG | Reverse RT-qPCR primer used to assess relative AdProt expression. |
| 5 | GTCAGGCACCGACTCCAACT | Forward RT-qPCR primer used to assess relative AcrIIA4–csd expression. |
| 6 | GTTCCAGCCGTTTTTAAATGCAC | Reverse RT-qPCR primer used to assess relative AcrIIA4–csd expression. |
| 7 | ACCCAGAAGGGACAGAAGAACAG | Forward RT-qPCR primer used to assess relative dCas9–TAD expression. |
| 8 | TTCTGCAGGTAGTACAGGTA | Reverse RT-qPCR primer used to assess relative dCas9–TAD expression. |
| 9 | CACCGACACCCCATTGACGTCAAT | Forward RT-qPCR primer used to assess relative expression of tiled gRNA #1. |
| 10 | CACCACGTCAGCTGCCAGATCCCA | Forward RT-qPCR primer used to assess relative expression of tiled gRNA #2. |
| 11 | CACCTGGGAGAACAGATCTGGCCT | Forward RT-qPCR primer used to assess relative expression of tiled gRNA #3. |
| 12 | CACCGCGCTAGCGAGCTCAGGTAC | Forward RT-qPCR primer used to assess relative expression of tiled gRNA #4. |
| 13 | CACCGAGAACAGATCTGGCCTCGG | Forward RT-qPCR primer used to assess relative expression of tiled gRNA #5. |
| 14 | CACCTGAAAGACGTCACAGTATGA | Forward RT-qPCR primer used to assess relative expression of tiled gRNA #6. |
| 15 | CACCAGTATGACGGCCATGGGATC | Forward RT-qPCR primer used to assess relative expression of tiled gRNA #7. |
| 16 | GACTCGGTGCCACTTTTTCAAG | Universal reverse primer used to assess relative expression of all gRNAs, anneals to the scaffold region of gRNAs. |
| 17 | GTCTCCACCGAGCTGAGAGA | Forward primer used to amplify dCas9 region of viral genome for sequencing. |
| 18 | ATGGACAAGAAGTACTCCATTGGGC | Reverse primer used to amplify dCas9 region of viral genome for sequencing. |
| 19 | TAATACGACTCACTATAGAC-GCATAAAGATGAGACGCGTTTTA-GAGCTAGAAAT | T7- and sgRNA spacer sequence containing forward primer for generating linear dsDNA templates for sgRNA synthesis (BLI) |
| 20 | AAAAGCACCGACTCGGTGCCAC-TTTTTCAAGTTGATAAC-GGACTAGCCTTATTTTAACTTGC-TATTTCTAGCTCTAAAAC | Universal dCas9 reverse primer for generating linear dsDNA templates for sgRNA synthesis (BLI, CASFISH assays) |

**Table S7 continued:**

| | | |
|---|---|---|
| 21 | Biotin-AGCAGAAATCTCTGCTGAC-GCATAAAGATGAGACGCTGGAG-TACAAACGTCAGCT | Top strand of the biotinylated dsDNA substrate containing the TGG PAM sequence (for BLI assay) |
| 22 | AGCTGACGTTT-GTACTCCAGCGTCTCATCTTTATGC GTCAGCAGAGATTTCTGCT | Bottom strand of the dsDNA substrate containing the PAM sequence (for BLI assay) |
| 23 | Biotin- AGCAGAAATCTCTGCTGAC-GCATAAAGATGAGACGCTCGAG-TACAAACGTCAGCT | Top strand of the biotinylated dsDNA substrate containing no PAM (TCG) sequence (for BLI assay) |
| 24 | AGCTGACGTTT-GTACTCGAGCGTCTCATCTTTATGC GTCAGCAGAGATTTCTGCT | Bottom strand of the dsDNA substrate containing no PAM sequence (for BLI assay) |
| 25 | TAATACGACTCACTATAC-CATATTCCACGTCCTACAGGTTTTA-GAGCTAGAAAT | T7- and sgRNA spacer sequence containing forward primer for generating linear dsDNA templates for sgRNA synthesis (CASFISH assay) |

### SUPPLEMENTARY FIGURES

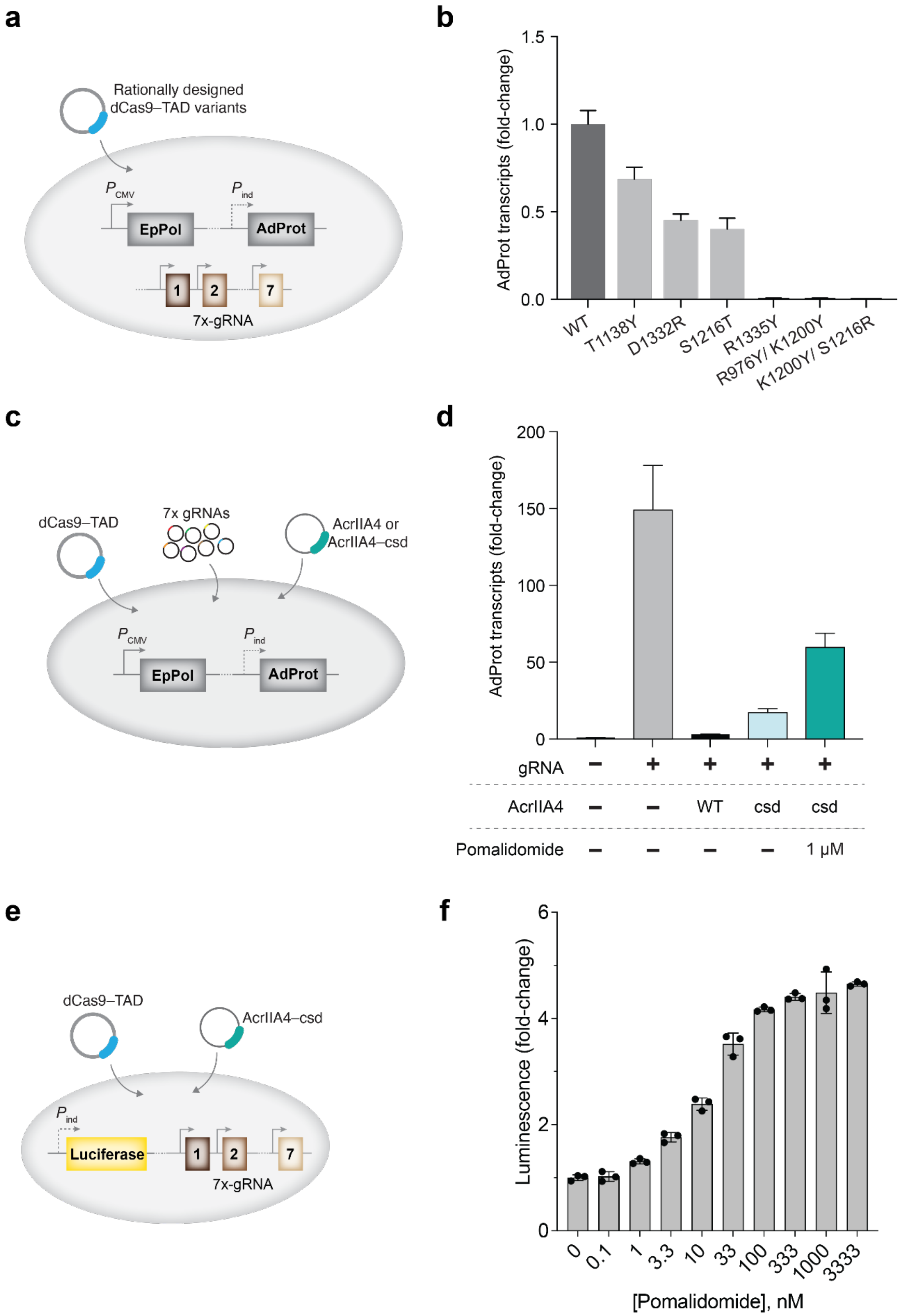

**Figure S1. (a)** Experimental setup to test previously reported[6] AcrIIA4-resistant dCas9 variants rationally designed to evade AcrIIA4 inhibition. dCas9–TAD variant plasmids were transfected into a cell line stably expressing tiling gRNAs targeting a promoter-less AdProt gene, and AdProt transcript levels were quantified by RT–qPCR. **(b)** All tested variants had reduced activity compared to wild-type dCas9–TAD, with some having no detectable activity. **(c)** Experimental setup to test whether AcrIIA4–csd enables pomalidomide-inducible relief of AcrIIA4-mediated inhibition. **(d)** In cells co-expressing dCas9–TAD, tiling gRNAs, and either AcrIIA4 or AcrIIA4–csd, AdProt transcript levels were restored only upon pomalidomide treatment in the AcrIIA4–csd condition, confirming small-molecule–dependent virus amplification. **(e)** Experimental setup to test whether pomalidomide can dosably control dCas9–TAD transcriptional activity. **(f)** Dose–response assay in a HEK293T luciferase reporter line co-transfected with dCas9–TAD and AcrIIA4–csd. Increasing pomalidomide concentrations led to graded restoration of CRISPRa-driven luminescence, demonstrating tunable control of dCas9–TAD activity via pomalidomide-mediated AcrIIA4–csd degradation.

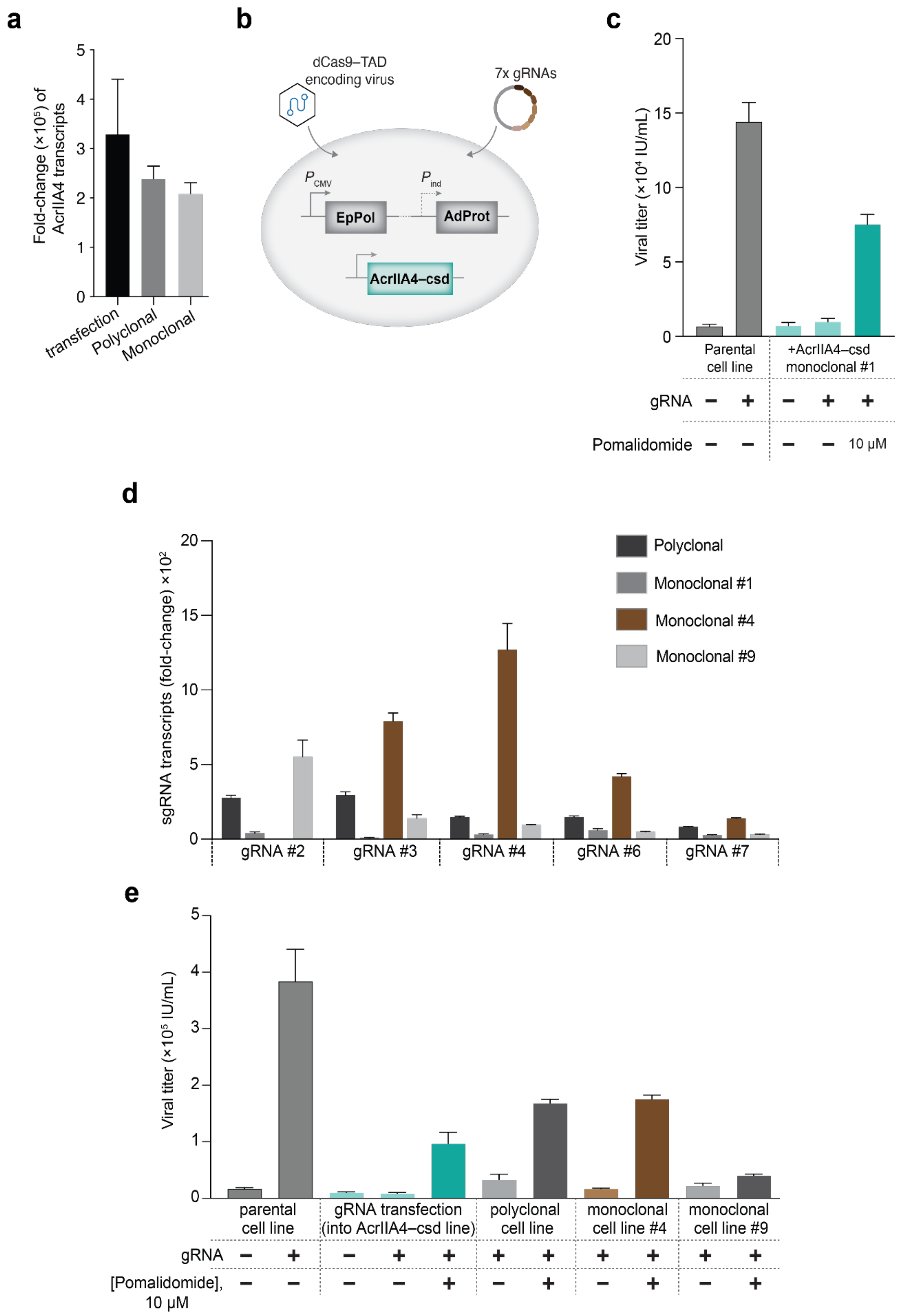
a
b
c
Fold-change (×10⁵) of AcrIIA4 transcripts
transfection
Polyclonal
Monoclonal
dCas9–TAD encoding virus
7x gRNAs
P CMV
P ind
EpPol
AdProt
AcrIIA4–csd
Viral titer (×10⁴ IU/mL)
Parental cell line
+AcrIIA4–csd monoclonal #1
gRNA
Pomalidomide
10 μM
d
sgRNA transcripts (fold-change) ×10²
Polyclonal
Monoclonal #1
Monoclonal #4
Monoclonal #9
gRNA #2
gRNA #3
gRNA #4
gRNA #6
gRNA #7
e
Viral titer (×10⁵ IU/mL)
parental cell line
gRNA transfection (into AcrIIA4–csd line)
polyclonal cell line
monoclonal cell line #4
monoclonal cell line #9
gRNA
[Pomalidomide], 10 μM

**Figure S2. Generation and validation of AcrIIA4–csd–expressing selector cell lines. (a)** qPCR analysis of AcrIIA4–csd transcript levels in transiently transfected, polyclonal, and monoclonal (clone #1) cell lines already stably expressing EpPol and promoter-less AdProt. **(b)** Schematic of viral replication assay performed in AcrIIA4–csd monoclonal line #1 to test pomalidomide responsiveness. **(c)** Viral replication in AcrIIA4–csd monoclonal cells was inhibited in the absence of pomalidomide. **(d)** qPCR analysis of gRNA expression in polyclonal and monoclonal selector lines stably encoding EpPol, AdProt (no promoter), and AcrIIA4–csd. **(e)** Pomalidomide-dependent rescue of viral replication in parental (no AcrIIA4–csd control), gRNA transfected, polyclonal, and two of the multiple screened monoclonal AcrIIA4–csd and gRNA expressing cell lines. Monoclonal #4 and the polyclonal line (both encoding EpPol, AdProt (no promoter), AcrIIA4–csd, and a gRNA cassette) were used for subsequent evolution experiments. For panels **c, d, e**: "Parental cell line" refers to the cells that do not express AcrIIA4. IU/mL = infectious units per mL, measured using flow cytometry.

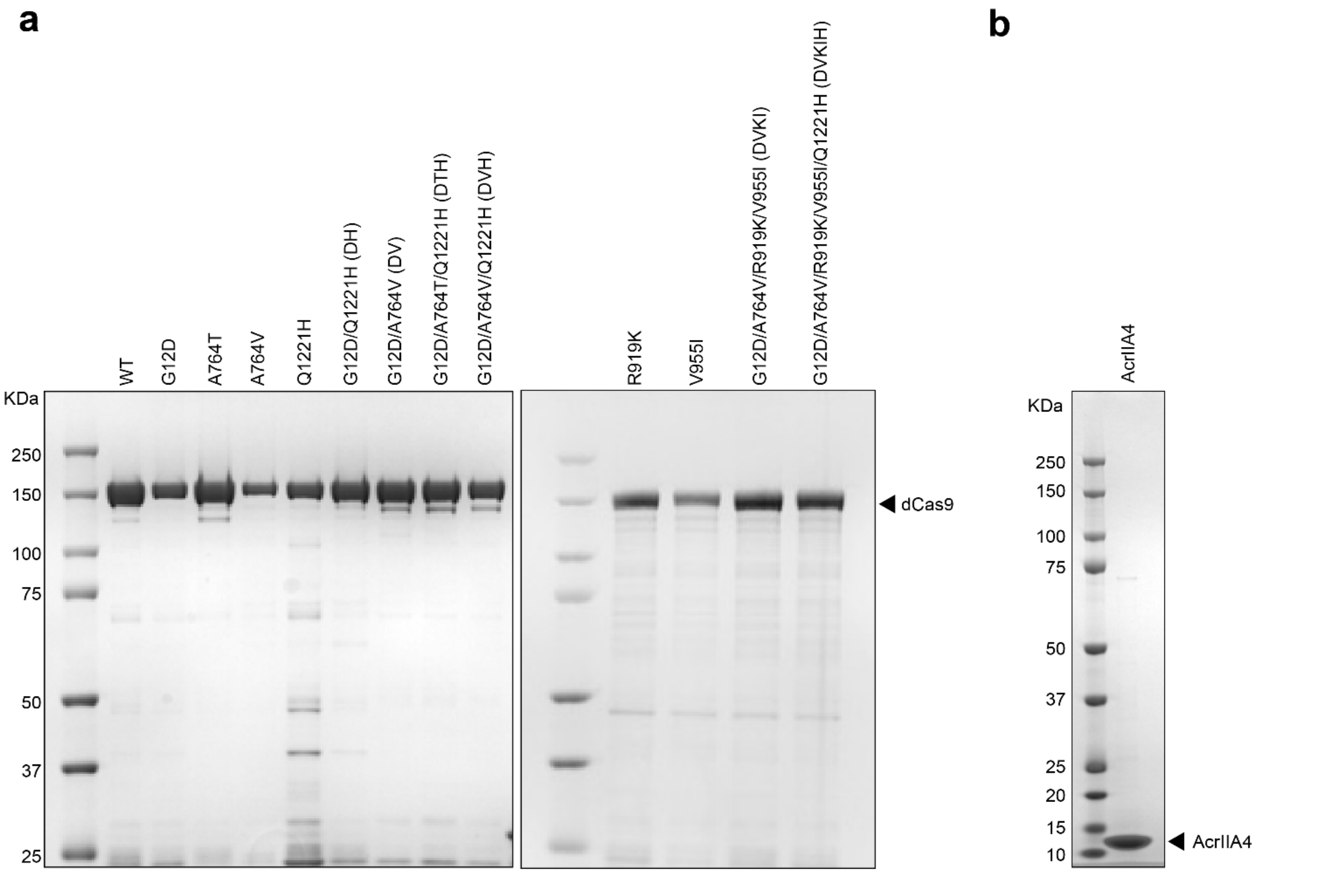


**Figure S3. (a–b)** SDS-PAGE gels for purified **(a)** dCas9 variants and **(b)** AcrIIA4 used in the biolayer interferometry experiments.

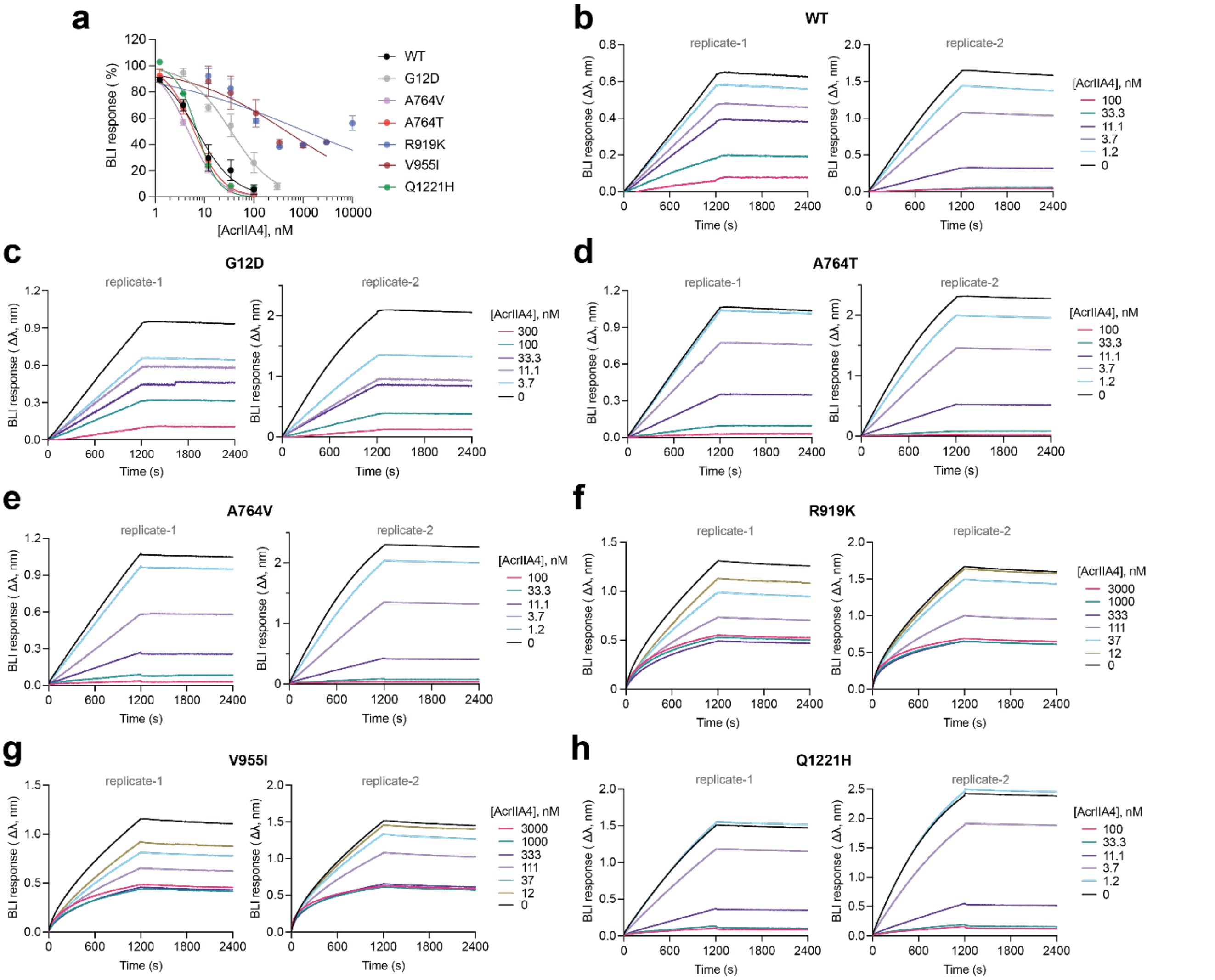

**Figure S4. BLI analysis of AcrIIA4-mediated inhibition of dCas9–DNA binding across singly-substituted dCas9 variants. (a)** Dose-dependent inhibition of dCas9–DNA binding by AcrIIA4 across wild-type (WT) dCas9 and indicated dCas9 variants, measured by normalized BLI response. **(b–h)** BLI sensograms showing inhibition of RNP–DNA binding at increasing AcrIIA4 concentrations for (**b**) WT, (**c**) G12D, (**d**) A764T, (**e**) A764V, (**f**) R919K, (**g**) V955I, and (**h**) Q1221H variants. Two biological replicates are shown for each variant.

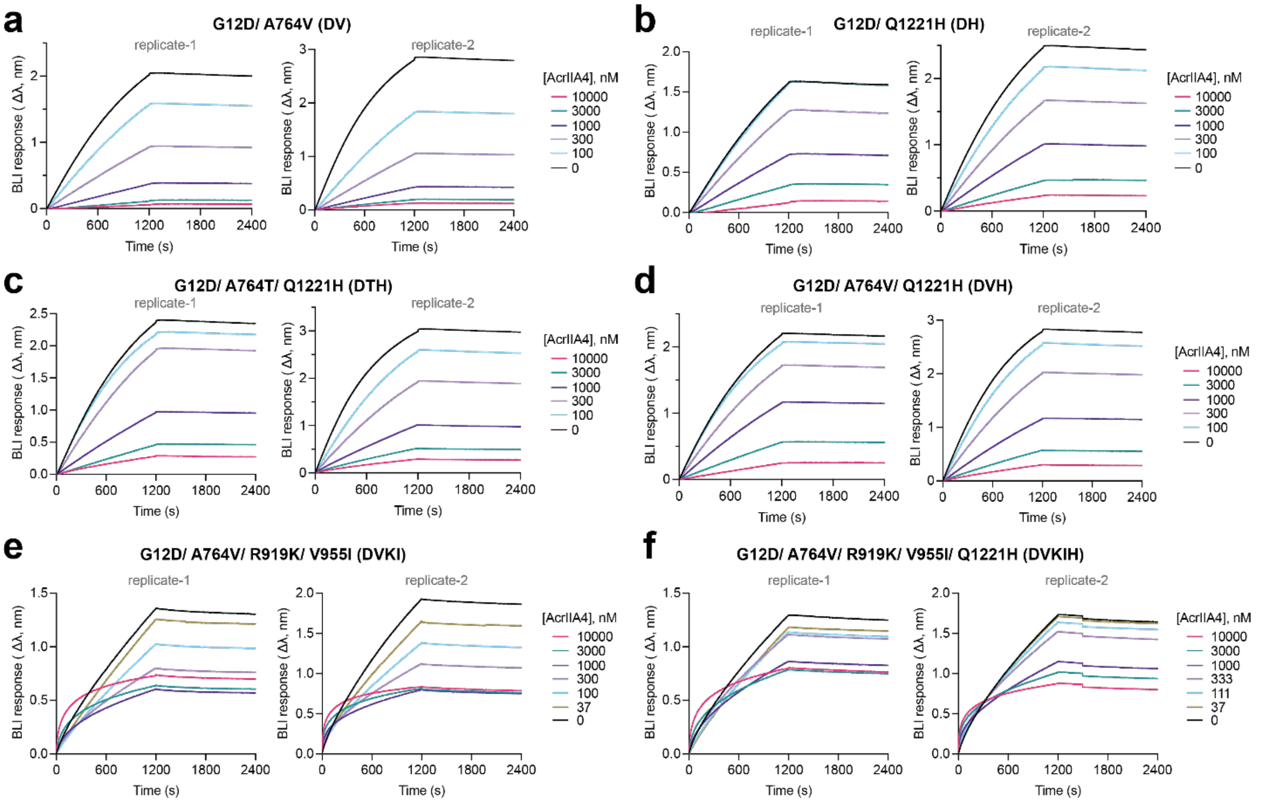


**Figure S5. BLI analysis of AcrIIA4-mediated inhibition of dCas9–DNA binding across evolutionarily relevant dCas9 variant RNPs. (a–f)** BLI sensograms showing binding of dCas9 RNP variants to PAM-containing DNA in the presence of increasing AcrIIA4 concentrations. Variants include (**a**) DV, (**b**) DH, (**c**) DTH, (**d**) DVH, (**e**) DVKI, and (**f**) DVKIH. Two biological replicates are shown for each variant.

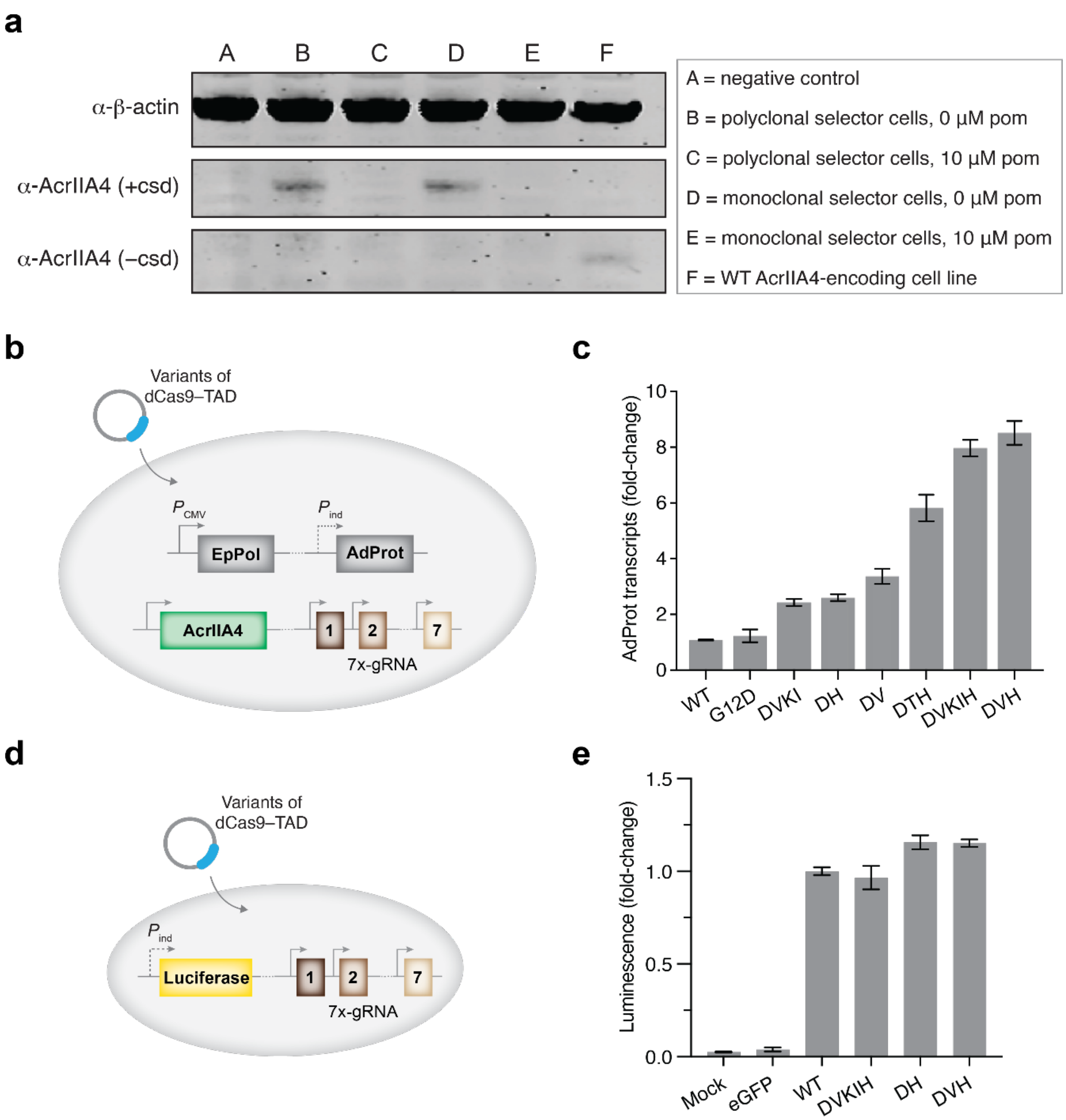


**Figure S6. (a)** Western blot analysis of AcrIIA4 expression using an α-AcrIIA4 antibody. Cell lines expressing AcrIIA4–csd or wild-type AcrIIA4 showed comparable AcrIIA4 expression levels. **(b)** A cell line stably expressing wild-type AcrIIA4, promoter-less AdProt, and gRNAs targeting the latter was used to assess transcriptional activation. **(c)** RT–qPCR analysis of AdProt expression following transient transfection of the indicated dCas9–TAD variants into the cell line from panel (**b**), evaluating their ability to activate AdProt transcription in the presence of wild-type AcrIIA4. **(d)** A HEK293T cell line encoding a promoter-less luciferase and stably expressing gRNAs targeting this region was used to assess the basal (no AcrIIA4) transcriptional activity of observed dCas9–TAD variants. **(e)** Results from luminescence assays conducted after the transfection of select, top-performing dCas9–TAD variants into the aforementioned cell line.

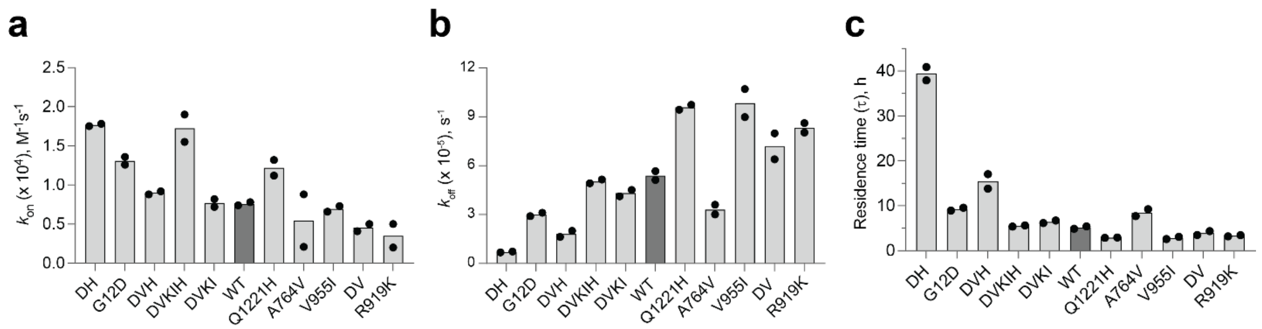


**Figure S7. Biolayer interferometry measurements of dCas9 RNP–DNA binding kinetics. (a–c)** Kinetic binding parameters derived from global fits of BLI sensograms for dCas9 variant RNP complexes with target DNA containing a PAM. Dissociation rate constants ($k_{off}$, **a**), association rate constants ($k_{on}$, **b**), residence times (τ) representing complex stability, calculated as $1/k_{off}$ (**c**). Each bar represents the mean of two replicates.

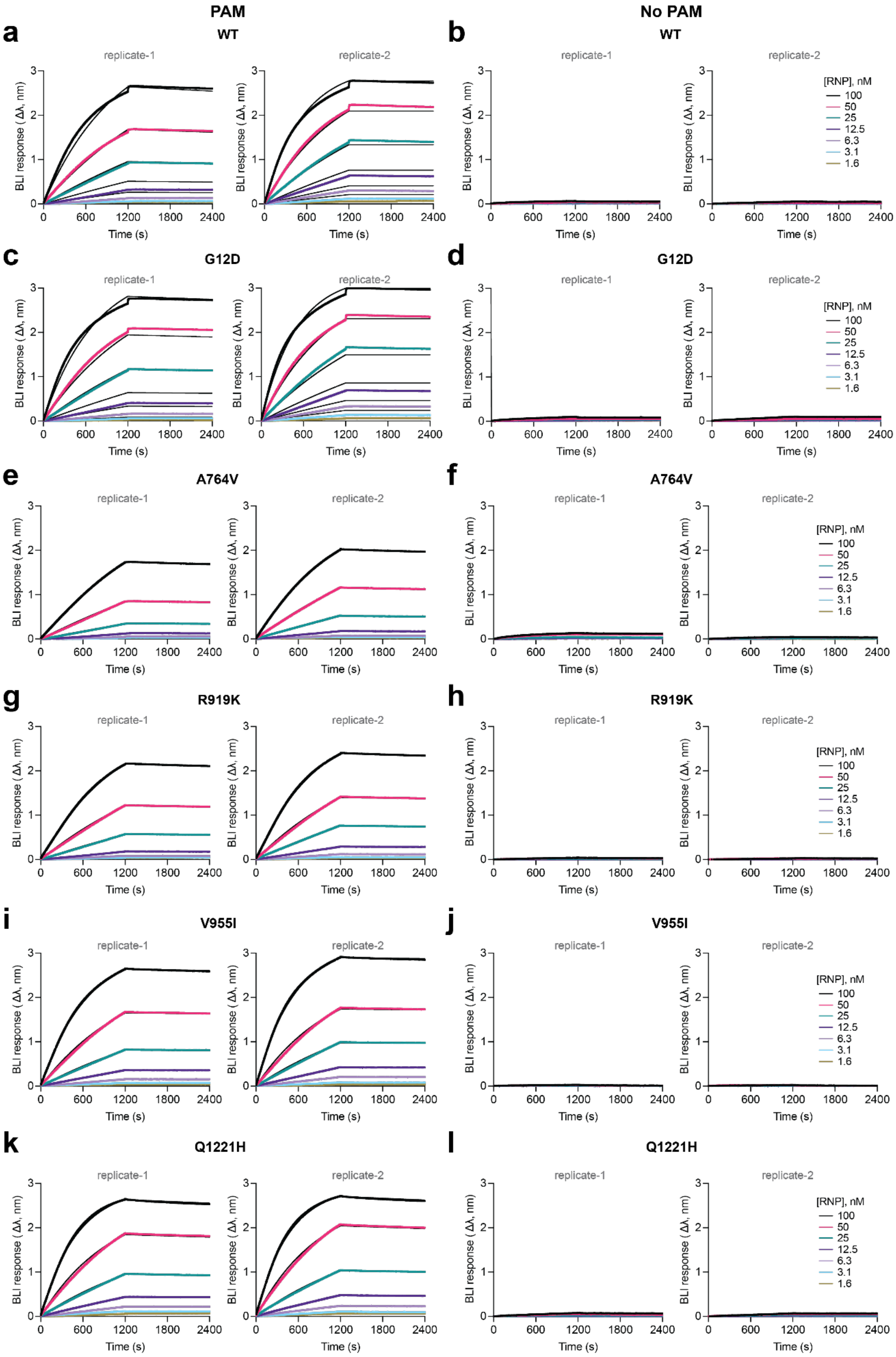
PAM
No PAM
a
b
WT
c
d
G12D
e
f
A764V
g
h
R919K
i
j
V955I
k
l
Q1221H
replicate-1
replicate-2
BLI response (Δλ, nm)
Time (s)
0
600
1200
1800
2400
[RNP], nM
100
50
25
12.5
6.3
3.1
1.6

**Figure S8. BLI sensograms of wild-type and singly-substituted dCas9 variant RNPs binding to PAM and no PAM DNA. (a–l)** BLI binding responses for wild-type (WT) dCas9 RNP and indicated dCas9 RNP variants at increasing RNP concentrations (1.6–100 nM) using PAM-containing (**a, c, e, g, i, k**) or PAM-lacking (**b, d, f, h, j, l**) DNA substrates. Robust, PAM-dependent binding was observed only for PAM-containing DNA. Two biological replicates are shown for (**a, b**) WT, (**c, d**) G12D, (**e, f**) A764V, (**g, h**) R919K, (**i, j**) V955I, and (**k, l**) Q1221H variants.

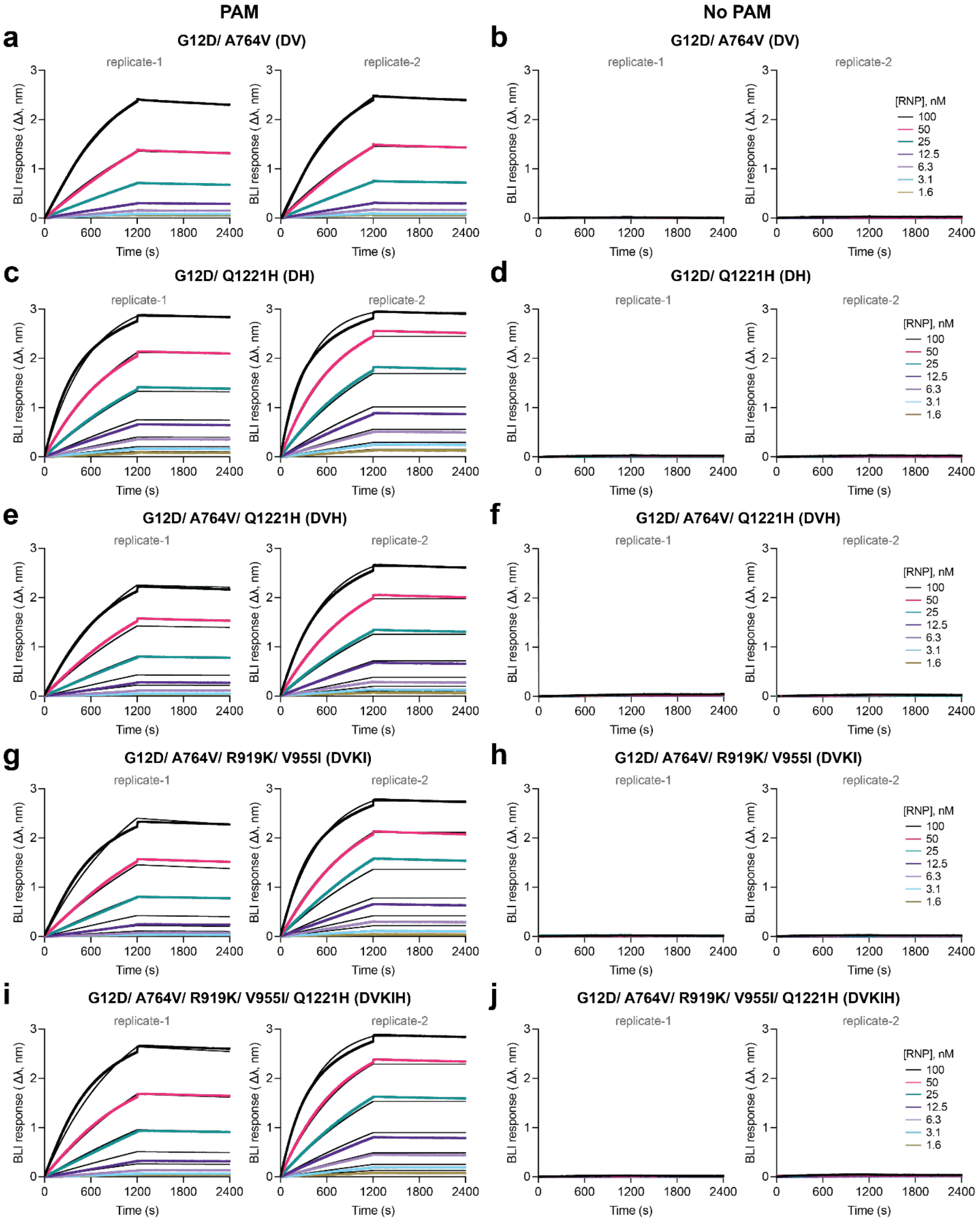


**Figure S9. BLI sensograms of evolutionary relevant dCas9 variant RNPs binding to PAM and no PAM DNA. (a–j)** BLI binding responses for indicated dCas9 RNP variants at increasing RNP concentrations (1.6–100 nM) using PAM-containing (**a, c, e, g, i**) or PAM-lacking (**b, d, f, h, j**) DNA substrates. Two biological replicates are shown for (**a, b**) DV, (**c, d**) DH, (**e,f**) DVH, (**g, h**) DVKI, and (**i, j**) DVKIH variants.

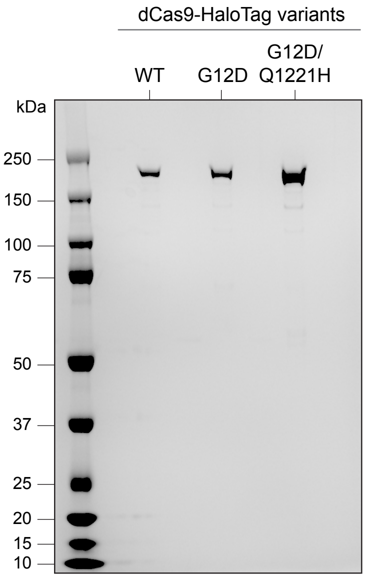


**Figure S10.** SDS-PAGE gel for purified dCas9-HaloTag variants used in the CASFISH experiment.

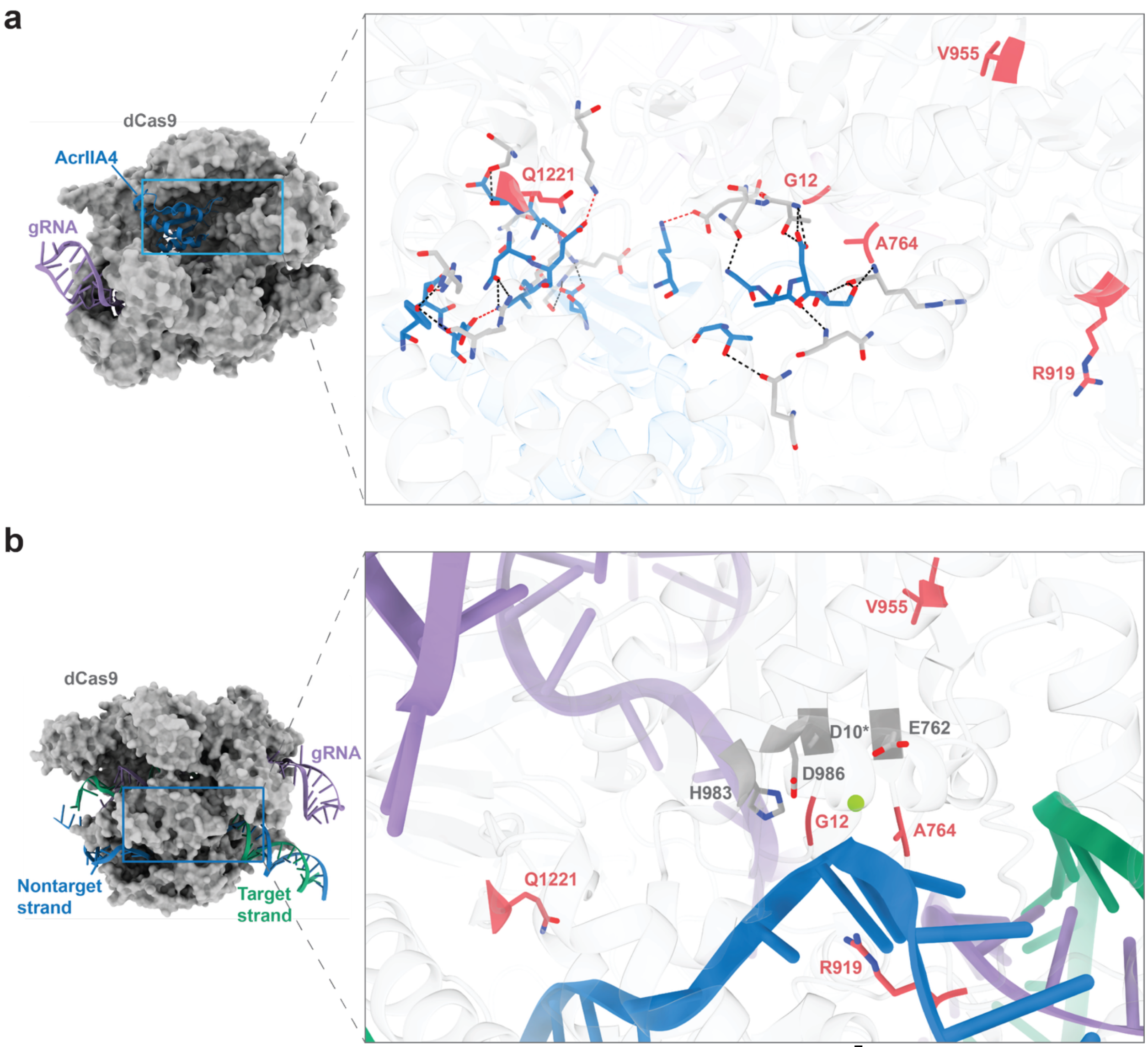


**Figure S11. (a)** The Cas9-gRNA-AcrIIA4 ternary complex (PDB 5VW1).[7] Cas9 (*grey*) is shown in surface representation, while AcrIIA4 (*blue*) and the gRNA (*purple*) are shown in cartoon representations. The inlet shows stick representations of the interaction interfaces between AcrIIA4 and Cas9 within the ternary complex. The Cas9 residues at which substitutions were identified are labeled and colored red. Hydrogen bonds and electrostatic interactions between Cas9 and AcrIIA4 are shown as black and red dashed lines, respectively. **(b)** The Cas9-gRNA-dsDNA ternary complex (PDB 5Y36).[8] Cas9 (*grey*) is shown in surface representation, while the gRNA (*purple*), the nontarget strand (*blue*), and the target strand (green) are shown in cartoon representations. The inlet shows stick representations of the RuvC domain with the nontarget strand (shown in cartoon representation). Residue D10 was substituted for alanine in the generation of this structure, as is present in catalytically inactive dCas9.
